# A type I 3’ UTR-derived sRNA is involved in heme metabolism and virulence in *Staphylococcus aureus*

**DOI:** 10.1101/2024.10.25.620293

**Authors:** Chloé Silard, Noëlla Germain-Amiot, Pierre Poirette, Julie Legros, Gabriella Duraõ, Sophie Martinais, Nicolas Mirouze, Yoann Augagneur

**Author notes:** Corresponding author at: Université de Rennes, BRM (Bacterial RNAs and Medicine) UMR_S 1230, Rennes, France.

## Abstract

*Staphylococcus aureus* is a pathogen responsible for a wide array of superficial to life-threatening infections. To efficiently adapt to environmental cues, a complex regulatory network is needed, involving among others regulatory RNAs (sRNAs). Here, we studied Srn_9342, an sRNA transcribed into two isoforms of different lengths and known to interact with RNAIII, leading to a modulation of δ-hemolysin production. We showed that the two isoforms are oppositely expressed in a growth phase-dependent manner. Then, we demonstrated using transcriptional fusions and various chromosomally recombinant strains that Srn_9342 is a type I 3’UTR-derived sRNA whose the expression of the long form (Srn_9342_L_) is SigB-dependent. Using a *Δsrn_9342* mutant, we monitored the transcript level of various RNA partners previously identified by MAPS and showed that the *hemQ* mRNA level, encoding a protein involved in heme biosynthesis and reported to participate in small colony variant (SCV) phenotype, increased in the mutant. *In silico* and *in vitro* biochemical investigations showed that a 5’ region of Srn_9342 binds *hemQ* leading to a repression of the HemQ protein level whereas the overexpression of Srn_9342 induced an SCV phenotype which was partially relieved by the addition of hemin. Finally, we report that the deletion of *srn_9342* significantly increased the virulence of the pathogen in a *Galleria mellonella* model. Taking together, these data uncovered a novel type I 3’ UTR-derived sRNA regulating the heme biosynthesis pathway and implicated in virulence and SCV formation in *S. aureus*.

## INTRODUCTION

*Staphylococcus aureus* is a major community- and hospital-acquired pathogen of humans and livestock. It is carried asymptomatically, either permanently or transiently, by around 30% of the population, most often in the nasal cavities. In certain conditions, it can cross the host barrier to initiate a wide spectrum of infections, ranging from superficial to severe. In addition, *S. aureus* is known to be resistant or recalcitrant to antibiotics therapy due to the acquisition of a mechanism of resistance, to its ability to enter in dormancy; or to exhibit a Small Colony Variant (SCV) phenotype [1–3]. Therefore, it possesses numerous regulatory systems to adapt to its environment and withstand attacks from the immune system during extra- or intra-cellular survival lifestyle. Among them, small regulatory RNAs (sRNAs) play an important role in numerus bacterial functions by acting as post-transcriptional modulators. They are usually defined as small, stable, generally non-coding RNAs with positive or negative regulatory functions on their targets. To date, several hundreds of sRNA candidates are identified in *S. aureus* [4], all compiled in the *Staphylococcus* regulatory RNA database (SRD) [5], with around 50 of them considered as *bona fide* [6].

The most studied staphylococcal sRNA is RNAIII, a paradigm of their role in bacterial physiology with its implication in the transition from commensalism to virulence [7, 8]. RNAIII is the effector of the *agrBDCA* system. It regulates a plethora of targets by binding to their mRNA, either as a repressor with or without the help of RNase III, but also sometimes as an activator [9–12]. Overall, it represses the expression of adhesion factors whereas it promotes the expression of several toxins. In *S. aureus*, other sRNAs such as SprD, SprC or RsaA, are known to be involved in virulence, either as pro- or as anti-virulent factors through their interaction with autolysin mRNA (*atl*), the *sbi* virulence factor mRNA, or *mgrA* mRNA encoding the global transcriptional regulator MgrA, respectively [13–15]. Since sRNAs are key players of virulence, they can be considered as attractive targets or active molecules to be used for the development of new treatments. Additionally, deciphering the mode of action of sRNAs could provide a better understanding of the regulatory network and virulence mechanisms of *S. aureus*, thereby aiding in its control.

As knowledge gained on sRNAs, they were classified into four different categories according to their mechanism of biogenesis: *cis*-antisense encoded sRNAs, *trans*-encoded sRNAs, or sRNAs derived from the 5’ or 3’ UTRs of mRNAs [16]. Whereas the first two categories are well-documented, examples of UTRs-derived sRNAs are scarcer. In *S. aureus*, Teg49 is an sRNA produced from the 5’ UTR of the SarA transcription factor mRNA after RNase III cleavage during the post-exponential phase of growth [17]. For their part, 3’ UTR-derived sRNAs have the particularity to be separated into two sub-categories. The Type I 3’ UTR-derived sRNAs, which are independent units of transcription possessing their own promoter within the CDS or 3’ UTR of an upstream gene; and the Type II 3’ UTR-derived sRNAs, which result from the cleavage of the 3’ UTR region of an mRNA by a specific RNase. Whereas there are some examples of Type II 3’ UTR-derived sRNAs in *S. aureus,* like RsaC and RsaG released from RNase III or RNase J1/J2 cleavage respectively [18, 19], there are no clear evidence of the existence of Type I 3’ UTR-derived sRNAs in this bacterial species. Srn_9342, an sRNA initially identified in Newman strain, is expressed from the 3’ end of a gene encoding a monofunctional amidase [20]. Therefore, it is falling in the 3’ UTR-derived sRNA category. A recent work showed that Srn_9342 is expressed under two forms of different lengths (Srn_9342_S_ and Srn_9342_L_), which are oppositely expressed as a function of growth phase [21]. Additionally, the use of MAPS to identified RNA targets of this sRNA [22] and revealed that Srn_9342_S_ preferentially bind sRNAs while Srn_9342_L_ has more affinity with mRNAs. Among sRNAs enriched during MAPS, RNAIII appeared as an attractive partner, with specific binding determined *in vitro*. Finally, it was shown that Srn_9342 slightly modulates the expression of the RNAIII-encoded δ-hemolysin during growth, suggesting a role in virulence [21].

Here, we further investigated the role of Srn_9342 in HG003, an NCTC_8325 derivative strain [23] in which the expression of RNAIII is more progressive due to a fully active SaeRS two-component system (TCS) [24, 25] and consider as a better model to study virulence and regulations in *S. aureus*. We first compared the *srn_9342* loci between the two strains and confirmed a common pattern of expression for the two Srn_9342 isoforms between the two strains. Then, we studied *srn_9342* promoter expression to decipher whether the sRNA is a type I or a type II 3’ UTR-derived sRNA using transcriptional fusions. In parallel, we monitored the expression of thirteen RNA targets identified by MAPS to unveil new role and function of Srn_9342. Among them, we showed that Srn_9342 repressed the expression of HemQ, a protein involved in heme biosynthesis and whose deletion is known to contribute to the formation of SCVs [26, 27], via a direct binding with *hemQ* mRNA. Thus, the overexpression of Srn_9342 led to the formation of tiny colonies whose phenotype could be partially relieved by the addition of hemin in extracellular environment. Finally, based on data collected in this study and on the interaction of Srn_9342 with RNAIII [21], we investigated the role of Srn_9342 in virulence using the *Galleria mellonella* model of infection and showed that its deletion significantly increased the virulence of *S. aureus*. Taken together, our data show that Srn_9342 is a new SigB-dependent type I 3’ UTR derived sRNA, which plays a role of an anti-virulent factor in *S. aureus*.

## RESULTS

### Srn_9342 expression is growth phase-dependent in HG003

Srn_9342 was previously identified in *S. aureus* Newman strain through the use of DETR’PROK workflow in order to identify novel non-coding transcripts from an RNAseq dataset [20, 28]. It is expressed under two forms of different lengths (Srn_9342_S_ and Srn_9342_L_) sharing the same 5’ end sequence. Using MAPS, it was shown that it can interact with various RNAs including mRNAs and sRNAs [21]. Among them, a specific interaction with RNAIII, leading to a modulation of δ-toxin production was demonstrated. In the present study, we aimed to identify the role of Srn_9342 in *S. aureus* using HG003, an NCTC_8325 derivative strain [29]. HG003 is a relevant model strain to study staphylococcal regulation and pathophysiology whereas Newman strain is known to be altered in *saeS*, which modify its phenotype compared to a more typical wild type *S. aureus* [24].

Since the *srn_9342* gene was found in the NM2 prophage of Newman, we compared the genetic organization of the *srn_9342* locus with HG003. In Newman, *srn_9342* overlaps with a 3’ end of a gene encoding an autolysin amidase domain (NWMN_1039) and is followed downstream by a cluster of two other sRNA genes (*srn_9344* and *srn_9345*) for which antisense genes *srn_9343* and *srn_9346* were also identified (**Fig. 1A**). In HG003, the cluster was not fully conserved as the two antisense *srn_9343*/*srn_9344 genes* were absent but replaced by three genes encoding two hypothetical proteins and one xenobiotic response element. However, the overlap between *srn_9342* and the autolysin amidase domain was retrieved (**Fig. 1A**). We performed an alignment of *srn_9342* and the 200 nucleotides (nts) upstream of the gene between the two strains (**Fig. 1B**). Overall, sequences share 99.55% of nts sequence identity, with only three single-nucleotide polymorphisms (SNP). Two of them were located in the putative promoter region at positions −200 and −59 nts and one in the sRNA sequence at +47 nts, which is outside the region shown to interact with RNAIII. This suggests a conservation in the mechanism of expression of Srn_9342 between the two strains.

**Figure 1:**
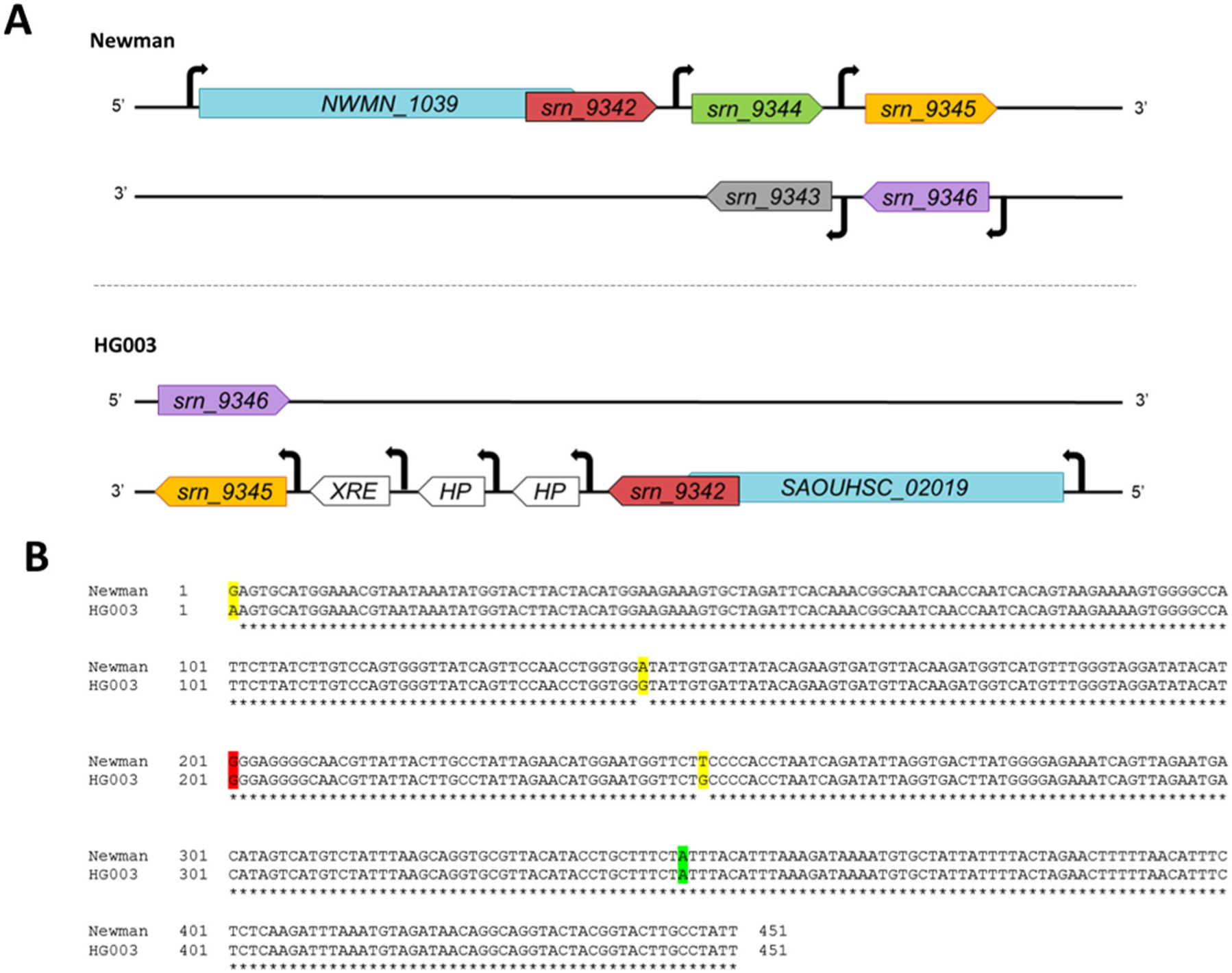
Comparison of the *srn_9342* loci between Newman and HG003 strains. (A) Genetic organization of the *srn_9342* locus in Newman strain (upper part) and HG003 strain (bottom part). (B) Alignment of *srn_9342* Newman and HG003 gene sequences including the 200 nts located upstream the 5’ end of the sRNA. Yellow: SNP; Red: 5’ end of Srn_9342_S/L_; Green: *srn_9342_S_* 3’ end.

To verify this, we studied Srn_9342 expression during growth in Tryptone Soya Broth (TSB). Total RNA was extracted every hour for 9 hours, then after 24 hours. The amount of transcript was monitored by Northern blot using a radiolabelled probe that hybridize the 5’ end of Srn_9342, allowing simultaneous detection of both isoforms (**Fig. 2A**). As for Newman [20], two distinct isoforms of Srn_9342 were detected: a short form (Srn_9342_S_) and a long form (Srn_9342_L_). The expression of each isoform depended on the growth phase, with Srn_9342_S_ being predominant in the early and exponential phases of growth, then being progressively substituted by Srn_9342_L_. After 24 hours of growth, Srn_9342_L_ was barely detected while Srn_9342_S_ was not (**Fig. 2A**). The proportion of each isoform was calculated, showing a switch of predominant form occurring after 5 hours of growth (OD_600nm_ of 5) where an equimolecular ratio was determined (**Fig. 2B**). The total amount of both Srn_9342_S_ and Srn_9342_L_ was also examined, revealing a nearly constant level of RNA transcripts over a 9-hour period (**Fig. S1**).

**Figure 2:**
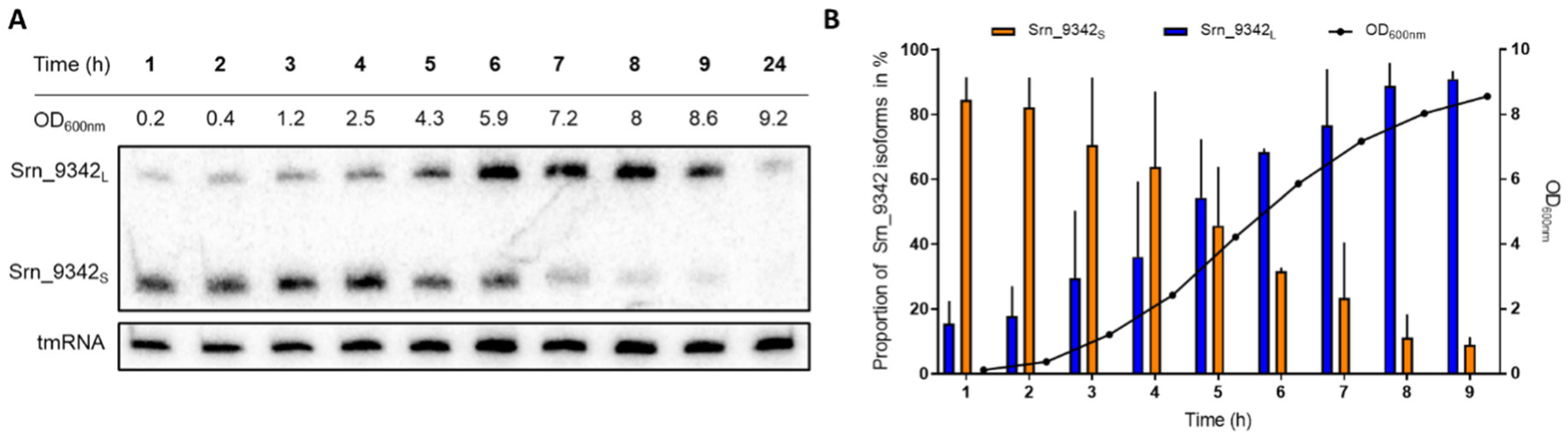
Srn_9342 isoforms expression in HG003 during growth. (A) Northern blot analysis of total RNA extracted each hour from *S. aureus* HG003 strain grown in TSB medium. A specific radioactive probe targeting the 5’ end of Srn_9342 was used, allowing the detection of both Srn_9342 isoforms simultaneously. tmRNA was used as an internal loading control. The Northern blot image is a representative experiment among three. (B) Quantification and proportion of each Srn_9342 isoform (Srn_9342_S_ = orange; Srn_9342_L_ = blue) over a 9-hour period of growth in TSB. The black line and points represent the OD_600nm_. Values are the mean of three independent biological replicates.

### Srn_9342 is a non-canonical type I 3’ UTR derived sRNA

To determine whether *srn_9342* has its own promoter, we cloned the putative promoter region of *srn_9342* into the plasmid pCN41c to create transcriptional fusions with the reporter gene *blaZ*, encoding β-lactamase, in HG003 [30]. To that aim, we tested different lengths of sequences (50 nts, 100 nts, and 200 nts) located upstream of the *srn_9342* 5’ end previously determined by RACE-PCR [21]. For each, we included the 6 nts downstream the 5’ end (yielding 57 nts, 107 nts, and 207 nts, respectively) and analysed the transcriptional activity against a promoterless control (0 nts). A transcriptional activity was detected for each promoter constructs whereas no reporter activity was detected for the control. The β-lactamase activity accumulated over growth with no significant differences between the different promoter lengths, indicating that *srn_9342* acts as an independent transcriptional unit and that all the elements essential for its expression are located within the first 50 nts upstream the transcriptional start site (TSS) (**Fig. 3A**).

**Figure 3:**
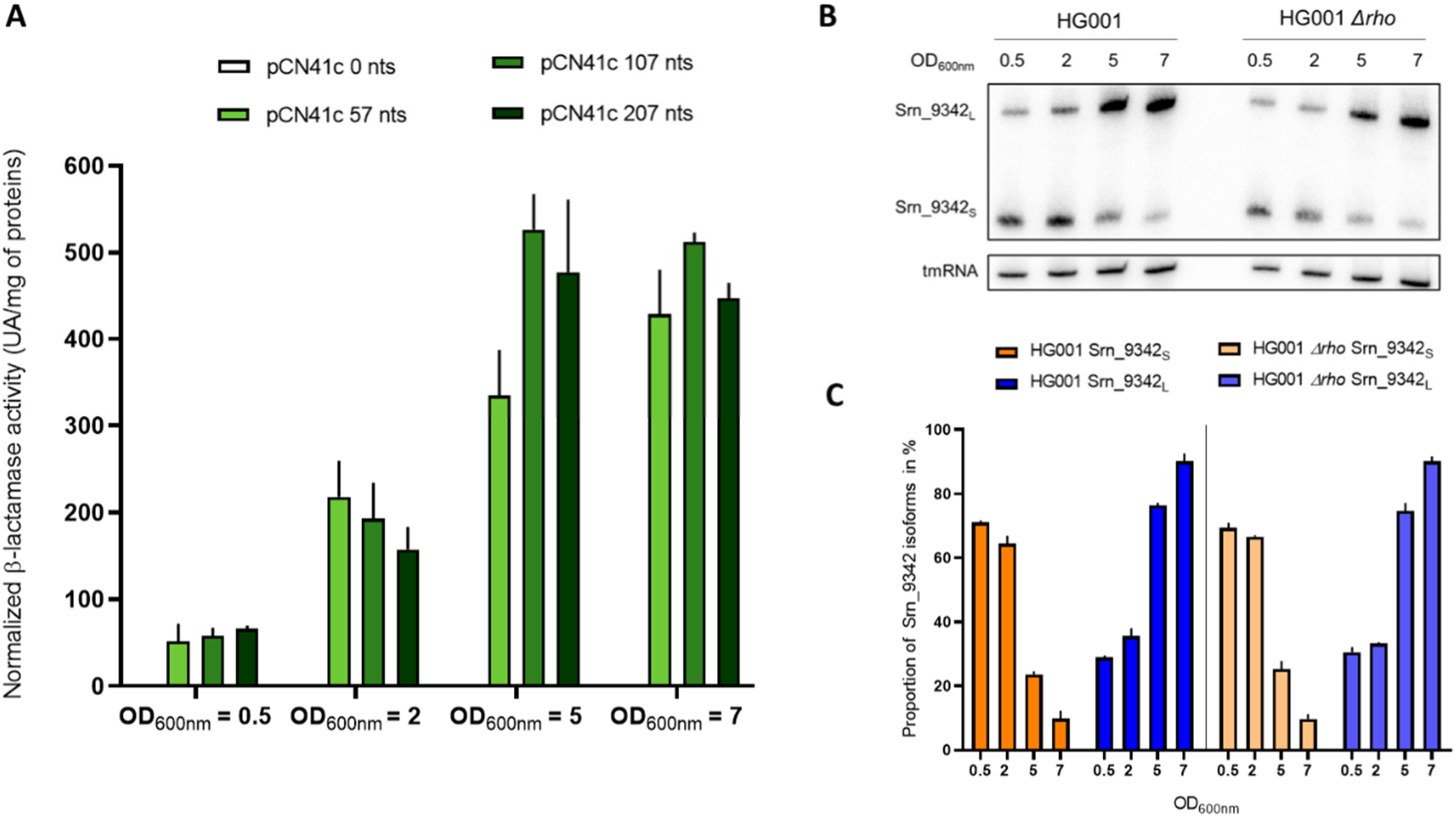
Srn_9342 is a non-canonical type I 3’ UTR-derived sRNA. (A) Detection of β-lactamase activity from transcriptional fusions of the *srn_9342* putative promoter with the *blaZ* reporter gene. Several lengths of 5’ upstream *srn_9342* sequences (0 nts, 57 nts, 107 nts, and 207 nts) were cloned in pCN41c and β-lactamase activity was detected at various time points of growth in HG003 strain. Values represent the mean of three independent experiments. (B) Northern blot analysis and (C) quantification of Srn_9342_S_ (orange) and Srn_9342_L_ (blue) isoforms during growth in HG001 (bright colours) and HG001 *Δrho* (pale colours). tmRNA was used as an internal loading control. The data presented are the mean of two Northern blot independent experiments.

Next, we investigated the mechanism of transcription termination of the *srn_9342* gene. To identify it, we compared the expression of Srn_9342 within HG001 or HG001 *Δrho* strains [31], which is defective for Rho protein responsible of Rho-dependant termination of transcription mechanism. HG001 strain is also derived from NCTC_8325 strain like HG003, but with *tcaR* gene not restored [23]. First, the pattern of expression of Srn_9342 was comparable between HG001 and HG003 (**Fig. 2A and 3B**). Second, there were no differences between the *Δrho* mutant and its parental HG001 strain in terms of expression profile and proportions of Srn_9342 isoforms, indicating that the transcription termination mechanism is not Rho-dependent (**Figure 3B and 3C**). Altogether, all these data indicate that Srn_9342 is expressed under two forms from an independent unit of transcription, and therefore is a type I 3’ UTR-derived sRNA.

### Srn_9342_L_ isoform expression is SigB dependent

We showed that Srn_9342 is expressed from an own promoter as is a type I 3’ UTR-derived sRNA. To further characterize its mechanism of expression we aimed to decipher whether a specific Sigma factor could be involved. Therefore, we searched for the presence of a Sigma factor binding sequence based on transcriptomic data from Mäder and colleagues in which Srn_9342 (named S810) transcript appeared as a 3’ UTR of SAOUHSC_02019 gene [31]. The authors identified a Sigma B (SigB) consensus (GGGTTAT-15-GGGTATTGT) located at −76 /-52 nts upstream of *srn_9342* 5’ end, which would be outside the 50 upstream nts sufficient for Srn_9342 expression (**Fig. 3A**). For this reason, we seeked for another SigB consensus within this sequence and identified a GTTACA-15nts-GGGTAG sequence (**Fig. 4A**), whose location at −10 / −35 nts is more appropriate to promote expression and shares similarity with the consensus used by Mäder (GGTTAA-15-GGGTAT), and other consensus published by Gertz and colleagues [32].

**Figure 4:**
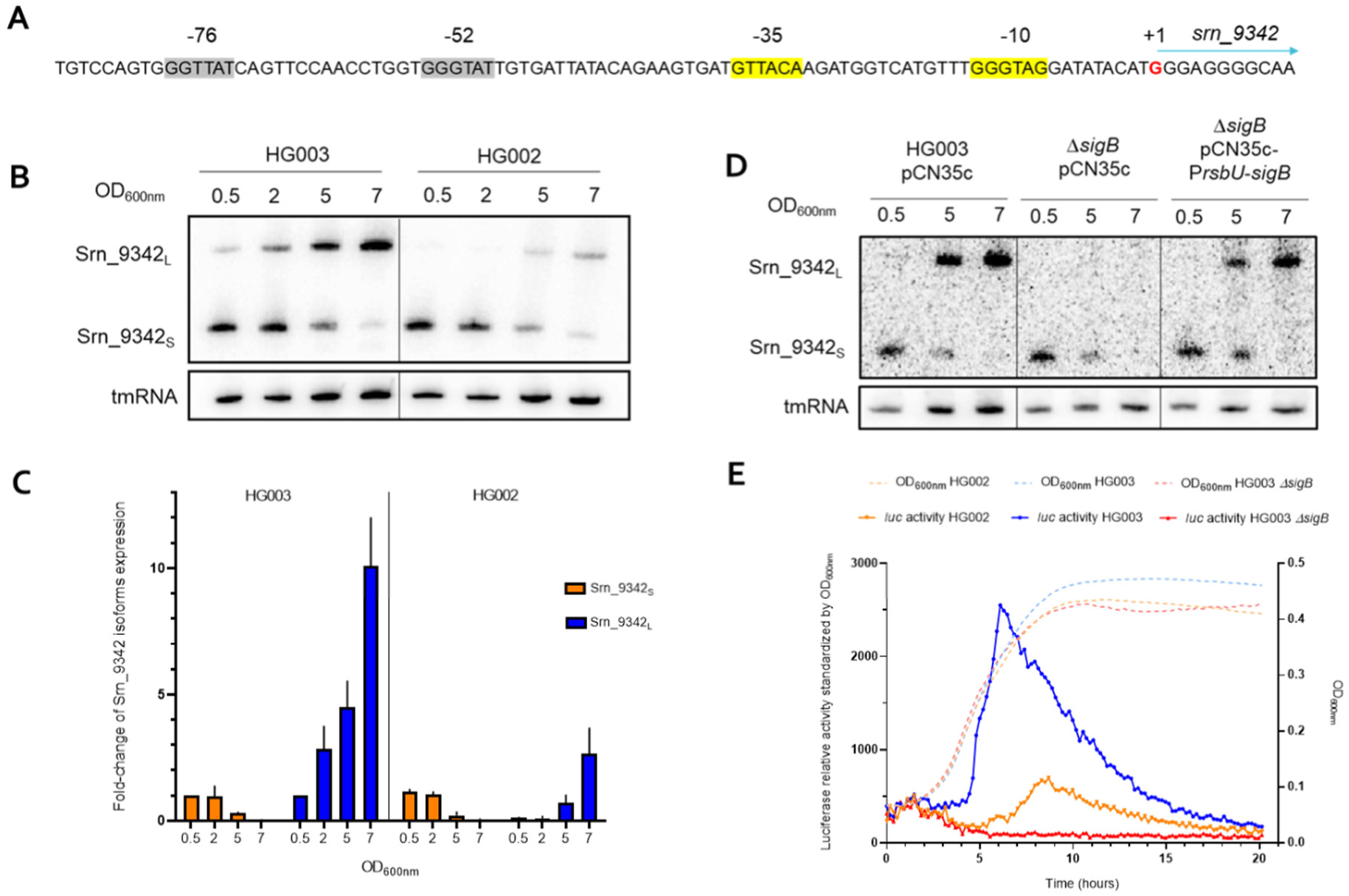
Srn_9342_L_ expression is SigB dependent. (A) Localization of putative SigB consensus sequences on *srn_9342* promoter. Grey: SigB consensus from Mäder *and al*. [31]; Yellow: SigB consensus found in this study; Red: TSS. (B) Expression of Srn_9342 and (C) determination of Srn_9342 isoforms expression in HG003 and HG002 during growth. Values of each isoform expression in HG003 at OD_600nm_ of 0.5 was adjusted to 1 and used as a standard. tmRNA was used as an internal loading control. The Northern blot image is a representative experiment among three independent replicates. (D) Expression of Srn_9342 isoforms in HG003 pCN35c, HG003 *ΔsigB* pCN35c, and HG003 *ΔsigB* pCN35c P*rsbU-sigB*. tmRNA was used as an internal loading control. The Northern blot image is a representative experiment among three independent replicates. (E) Real-time monitoring of luciferase activity of the 207 nts *srn_9342* promoter in HG003, HG002, and HG003 *ΔsigB* over 20 hours of growth. The luciferase activity was standardized by the OD_600nm_ measured every 10 minutes. Values are a mean of five (HG003 and HG002) or four (HG003 *ΔsigB*) biological replicates.

To confirm the involvement of the SigB factor in Srn_9342 expression, we compared the expression of Srn_9342 between HG003 and its isogenic mutant HG002, in which SigB is shut down due to a 11 nts deletion in the sequence of *rsbU* encoding the SigB positive regulator RsbU. Interestingly, Srn_9342_L_ was not detected at all at OD_600nm_ of 0.5 and 2 in HG002, whereas its expression level was significantly lower at OD_600nm_ of 5 and 7 compared to the strain HG003 (**Fig. 4B and 4C**). Conversely, the Srn_9342_S_ RNA level was not modified between the two strains in terms of pattern of expression and transcript quantity.

To further demonstrate the role of SigB in Srn_9342 expression, we used an HG003 *ΔsigB* mutant obtained by phage transduction of a *S. aureus* WCUH29 *ΔsigB*::Tc mutant in which *sigB* is replaced by a tetracycline resistance cassette [33]. Northern blot analysis showed that Srn_9342_L_ was totally absent when the SigB factor cannot be expressed, and its expression was fully restored when *sigB* is expressed from the pCN35c plasmid vector under the control of *rsbU* promoter, confirming the role of SigB in Srn_9342_L_ expression (**Fig. 4D and S2**).

Based on these new findings, we studied the expression of Srn_9342 in real-time using transcriptional fusion with *luc* reporter gene, encoding a firefly luciferase. Bioluminescence was then monitored during bacterial growth in HG003, HG002, and HG003 *ΔsigB* strains. As for the β-lactamase activity (**Fig. 3A**), reporter gene activity increased in post-exponential phase of growth, following Srn_9342_L_ transcript production (**Fig. 4E**). Conversely, the expression of firefly luciferase was drastically reduced in HG002 and completely abolished in HG003 *ΔsigB*, confirming the role of SigB for optimal expression *srn_9342* and that Srn_9342_L_ is SigB-dependent, but nor Srn_9342_S_.

### *srn_9342_L_* is necessary for Srn_9342_S_ expression

To further study the role of Srn_9342 in *S. aureus* physiology, we constructed a HG003 *Δsrn_9342* strain by chromosomic double-recombination. This *Δsrn_9342* mutant consist in the deletion of the whole *srn_9342* gene including its promoter (200 nts upstream the TSS). The deletion was verified by Northern blot with a complete absence of detection of each isoform (**Fig. 5A**). In addition, RT-qPCR analysis was performed to verify that the deletion of *srn_9342* do not lead to any polar effect on *SAOUHSC_02018* downstream gene (**Fig. S3**). Then, to discriminate the effects linked to one or other of the isoforms, we attempted to construct two chromosomally complemented strains derived from HG003 *Δsrn_9342* by cloning either the full *srn_9342* gene or the *srn_9342_S_* truncated gene under the control of the endogenous promoter (**Fig. S4**). Double recombination was done at the intergenic neutral locus SAPhB1502, as described by Coronel-Tellez and colleagues [34].

**Figure 5:**
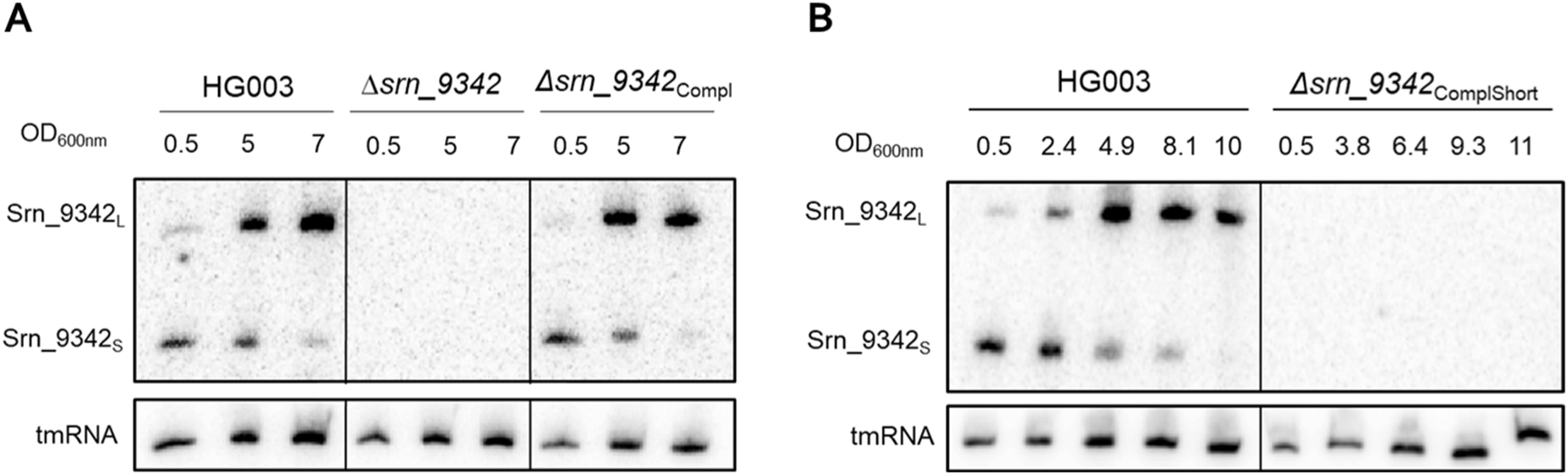
Full-length *srn_9342* gene is necessary for Srn_9342_S_ expression. (A) Expression of Srn_9342_S_ and Srn_9342_L_ in HG003, *Δsrn_9342* and *Δsrn_9342_Compl_*. tmRNA was used as an internal loading control. The Northern blot image is a representative experiment among three independent replicates. (B) Expression of Srn_9342_S_ and Srn_9342_L_ levels in HG003 and *Δsrn_9342_ComplShort_*. tmRNA was used as an internal loading control. The Northern blot image is a representative experiment among two independent replicates.

The strain complemented with the entire *srn_9342* gene (*Δsrn_9342*_Compl_) fully restored the Srn_9342 specific pattern of expression with the transition from Srn_9342_S_ to Srn_9342_L_, further confirming that Srn_9342 is a type I 3’ UTR-derived sRNA and that the transcription of the upstream gene (*SAOUHSC_02019*) does not play a role in *srn_9342* expression (**Fig. 5A and S5**). Conversely, no expression was detected in *Δsrn_9342*_ComplShort_ strain, despite the chromosomic insertion of all putative genetic information (endogenous promoter and putative Rho-independent terminator) necessary for Srn_9342_S_ expression, suggesting that the full version of the gene (including its 3’ end) is required for full expression of both Srn_9342 isoforms (**Fig. 5B**).

### Srn_9342 is a small regulatory RNA targeting *hemQ* mRNA and repressing HemQ production

Previous work on Newman strain led to the identification of a set of sRNAs and mRNAs that are potentially regulated by Srn_9342_S/L_ [21]. To unveil the most relevant targets, we analyzed by RT-qPCR the relative RNA level of thirteen of these targets (five sRNAs and seven mRNAs) in HG003 and *Δsrn_9342* at three time-points of growth (OD_600nm_ of 0.5 or 5 or 7) (**Fig. S6**). Among sRNAs identified by MAPS, we monitored the transcript level of the most enriched sRNAs (Sau6477, RNAIII, SprD, RsaC and VigR). Only RNAIII transcript level was modified in the *Δsrn_9342* mutant with a two-fold decrease at OD_600nm_ of 0.5, whereas no significant differences were detected later. Regarding mRNA targets, no significant variations were detected for *cibB*, *fbaA*, *recA* and *sirC* mRNAs. Conversely, significant differences were measured for *hemQ, SAOUHSC_00699* and *mvaS* mRNAs. The mRNA level of *hemQ* decreased by 2-fold at OD_600nm_ of 0.5 in the *Δsrn_9342* mutant, but then increased by 8-fold at OD_600nm_ of 7.5. Similarly, a 2-fold decreased was measured for *SAOUHSC_00699*, followed by an around 3-fold increase at OD_600nm_ of 7.5. Then, a 3.5-fold increase in the *mvaS* mRNA level was detected at OD_600nm_ of 7.5.

Based on this first RT-qPCR screen, we decided to focus exclusively on *hemQ* and verified that the transcript variations was actually due to Srn_9342 absence. To do so, we extracted RNAs at OD_600nm_ of 7 from the parental strain, the *Δsrn_9342* mutant and the *Δsrn_9342_Compl_* strain (**Fig. 6A**). In this second screen, we confirmed a 5-fold increase of *hemQ* in the *Δsrn_9342* strain which came back to the parental level in the *Δsrn_9342_Compl_* strain. These data indicate that Srn_9342_L_ negatively regulates *hemQ* mRNA level. To assess whether the Srn_9342_L_ amount is sufficient to support this regulation, we determined the relative levels of the two molecules in the HG003 strain at OD_600nm_ of 7 and found that there is fifty times more Srn_9342_L_ RNA compared to *hemQ* mRNA at this growth stage (**Fig. 6B**).

**Figure 6:**
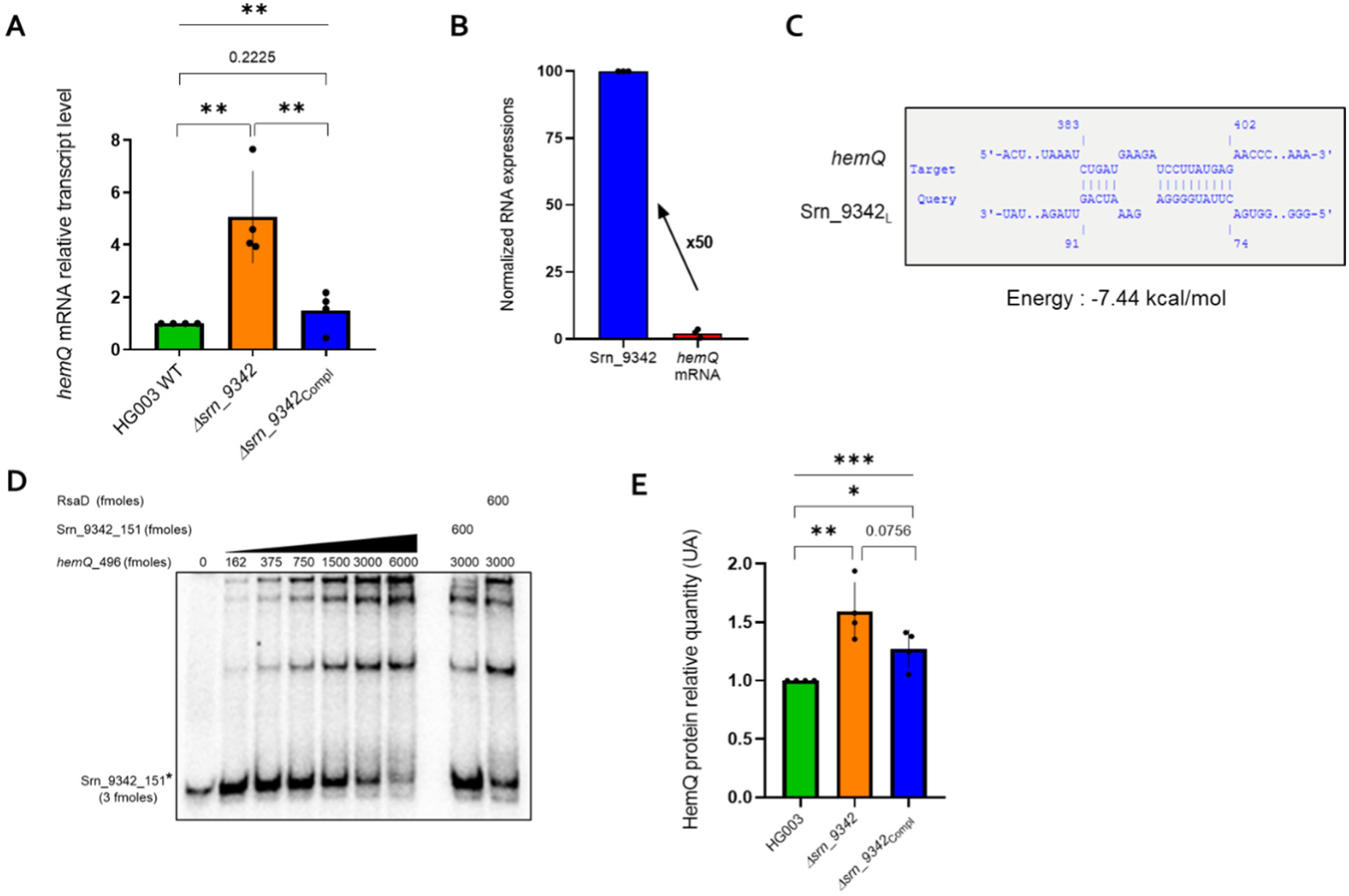
Srn_9342_L_ represses the expression of HemQ by binding onto its mRNA. (A) Effect of *srn_9342* gene deletion on the *hemQ* mRNA transcript level. Relative expression levels of *hemQ* mRNA determined by RT-qPCR on total RNA extracted from TSB cultures of HG003, *Δsrn_9342* and *Δsrn_9342_Compl_* at OD_600nm_ of 7. Values are the mean of four biological replicates. Data were analysed using the ΔΔCT method, with *gyrB* as an internal control and the HG003 strain as the calibrator. A Kruskal-Wallis non-parametric test across all conditions, followed by an unpaired t-test between pairs of data, was performed to determine significant differences among conditions (* p < 0.05, ** p < 0.01). (B) Proportion of *hemQ* mRNA quantity compared to Srn_9342_L_ in HG003 strain at OD_600nm_ of 7. Values are the mean of three biological replicates. Data were analysed using the ΔΔCT method, with *gyrB* as an internal control and the HG003 strain as the calibrator. (C) *In silico* prediction of interaction between *hemQ* mRNA and Srn_9342_L_ using IntaRNA. (D) Complex formation between unlabelled *hemQ* mRNA (496 nts) and radiolabelled *Srn_9342\** (151 nts). To assess complex formation specificity, an excess of unlabelled Srn_9342-151nts or RsaD sRNA were used. (E) Srn_9342 modulates HemQ expression. The amount of HemQ protein was quantified by Western blot in HG003, *Δsrn_9342* and *Δsrn_9342*_Compl_ level at OD_600nm_ of 7. Values represent the mean of four biological replicates. A Kruskal-Wallis non-parametric test across all conditions followed by unpaired t-test between pairs of data was performed to determine significant differences among conditions (* p < 0.05, ** p < 0.01, *** p < 0.001).

In addition, we performed *in silico* predictions of sRNA-mRNA interactions using the IntaRNA program [35]. IntaRNA identified a putative interaction between Srn_9342_L_ and *hemQ* mRNA with an energy of −7.44 kcal/mol (**Fig. 6C**). Based on this prediction, the 5’ moiety of Srn_9342_L_ (from nts 74 to 91) would interact with the coding sequence of *hemQ* (from nts 383 to 402). To test whether a direct interaction occurs between Srn_9342_L_ and *hemQ* mRNA, we performed an electrophoretic mobility shift assay (EMSA). An Srn_9342 fragment of 151 nts and a *hemQ* RNA fragment of 496 nts, both containing the predicted interaction regions, were *in vitro* transcribed. When increasing amounts of unlabelled *hemQ-*496nts mRNA were added to radiolabelled Srn_9342*, a specific complex was formed between the two RNAs (**Fig. 6D**). Indeed, the addition of RsaD sRNA did not disturb the complex using 3000 fmoles of *hemQ*, whereas an excess of unlabelled Srn_9342 displaced the complex.

To further confirm that Srn_9342 is a regulator of HemQ expression, we chromosomally inserted a FLAG sequence at 3’ end of *hemQ* and performed Western blot using an anti-FLAG antibody (**Fig. 6E**). Consistent with RT-qPCR measurements, a 50% increase in the HemQ protein level was detected at on OD_600nm_ of 7 in the *Δsrn_9342* strain, whereas it decreased to a level close to the parental in the complemented strain. Taken together, these data indicate that Srn_9342_L_ represses the expression of HemQ by binding onto its mRNA.

### Srn_9342 sRNA is linked to the SCV formation

Our data indicate that Srn_9342_L_ regulates the expression of HemQ, encoding a Coproheme decarboxylase involved in hemin biosynthesis [36]. In addition, a *hemQ* gene mutation is known to implicated in the development of the small colony variants (SCV) since its deletion produces tiny colonies which can be rescued by the addition of hemin in culture medium [26]. Since we showed that Srn_9342_L_ is a repressor of HemQ expression, we hypothesized that the overexpression of Srn_9342_L_ should affect HemQ expression and the overall fitness of the clone. To verify this, we cloned *srn_9342_S_* and *srn_9342_L_* genes under the control of P*srn_9342* or P*ami* promoters in the plasmid pCN38 [29]. Cloning of the full-length *srn_9342* gene allowed overexpression of both isoforms although the ratio between the two isoforms was modified (**Fig. S7**). Consistent with data obtained at the chromosomal level (**Fig. 5B**), cloning of Srn_9342_S_ under its own promoter did not lead to any expression of the sRNA whereas cloning under the P*ami* promoter was not viable.

When isolated on tryptic soy agar plates, overexpression strain of Srn_9342 resulted in a growth deficiency similar that an SCV phenotype compared with the parental strain and the *Δsrn_9342* mutant (**Fig. 7A**). The colonies were much smaller, and they displayed a significant growth delay in tryptic soy broth with a final biomass similar than that of the control strains. Interestingly, this phenomenon was repeatable as dilution of the overnight culture led to similar growth defect, suggesting that the phenotype was stable (**Fig. 7B**).

**Figure 7:**
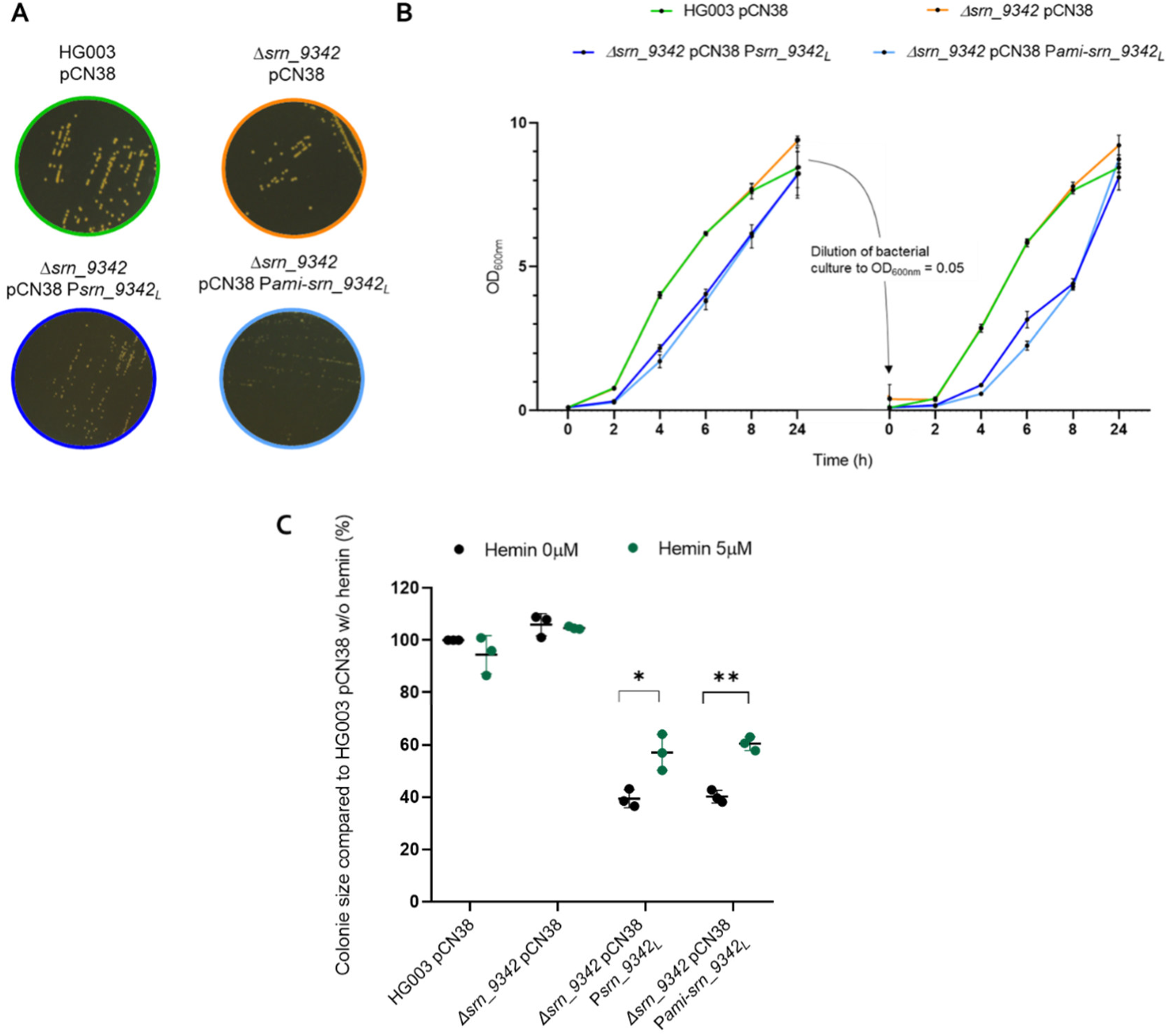
Srn_9342 overexpression induces an SCV phenotype partially restored by the presence of extracellular hemin. (A) Colony sizes on TSA plates with chloramphenicol (10 µg/mL), and (B) growth curves in TSB media with chloramphenicol (10 µg/mL) of HG003 and *Δsrn_9342* strains containing pCN38 empty-plasmid, compared to *Δsrn_9342* overexpression strains, containing the multi-copy plasmid with *srn_9342_L_* under its endogenous promoter (*Δsrn_9342* pCN38 P*srn_9342_L_*) or the P*ami* constitutive promoter (*Δsrn_9342* pCN38 P*ami-srn_9342_L_*). Values are the mean of three biological replicates. (C) Colony sizes on TSA plates, with (5 µM) or without (0 µM) hemin supplementation, expressed as a percentage relative to the parental strain without hemin control condition (left). Each condition represents a mean of three independent experiments, each containing ten colony size measurements. A two-way ANOVA followed by multiple t-tests was performed on the data (*: p-value < 0.05; **: p-value < 0.01).

Then, we verified whether the addition of hemin could restore the growth defect linked with the overexpression of Srn_9342. To that aim, strains were spotted on agar plates supplemented or not with 5 µM of hemin and colony size measurements were performed after 24 hours (**Fig. 7C**). Overall, a partial restoration of colony size was observed with a 20 % increase upon hemin addition. Conversely, the addition of hemin did not affect the colony size of the parental strain and its *Δsrn_9342* mutant (**Fig. 7C**). Therefore, these results indicate that Srn_9342 likely affects heme biosynthesis in *S. aureus* and contributes to the SCV phenotype through the regulation of HemQ protein production.

### Srn_9342 plays a role in *S. aureus* virulence

Based on the MAPS study showing that Srn_9342 can interact with various actors involved in virulence including RNAIII [21] and the role of Srn_9342 in the regulation of heme biosynthesis, we hypothesized that Srn_9342 may play a role in *S. aureus* pathogenicity. To verify this, we used the *Galleria mellonella* larvae model of infection and monitored the survival of the host during 6 days after injection of HG003, *Δsrn_9342* or *Δsrn_9342_Compl_* strains. Whereas infections with HG003 or Srn_9342_S+L_ complemented strains led to 64.8% and 64.2% of mortality respectively, more than 87% of the larvae were killed by the *Δsrn_9342* mutant (**Fig. 8A**). To verify whether this result was statistically significant, inoculum counts were performed for each strain in parallel to larvae infection (**Fig. 8B**). No standardization issues were observed with an actual inoculum found between 1.5 and 5 x 10^6^ CFU/mL, in accordance with a prior model standardization study [37]. Altogether, these findings confirm the implication of Srn_9342 in the virulence of *S. aureus* HG003 in an invertebrate infection model. Srn_9342 appears to act as an anti-virulence factor, like other *S. aureus* sRNAs such as SprC and RsaA [14, 15].

**Figure 8:**
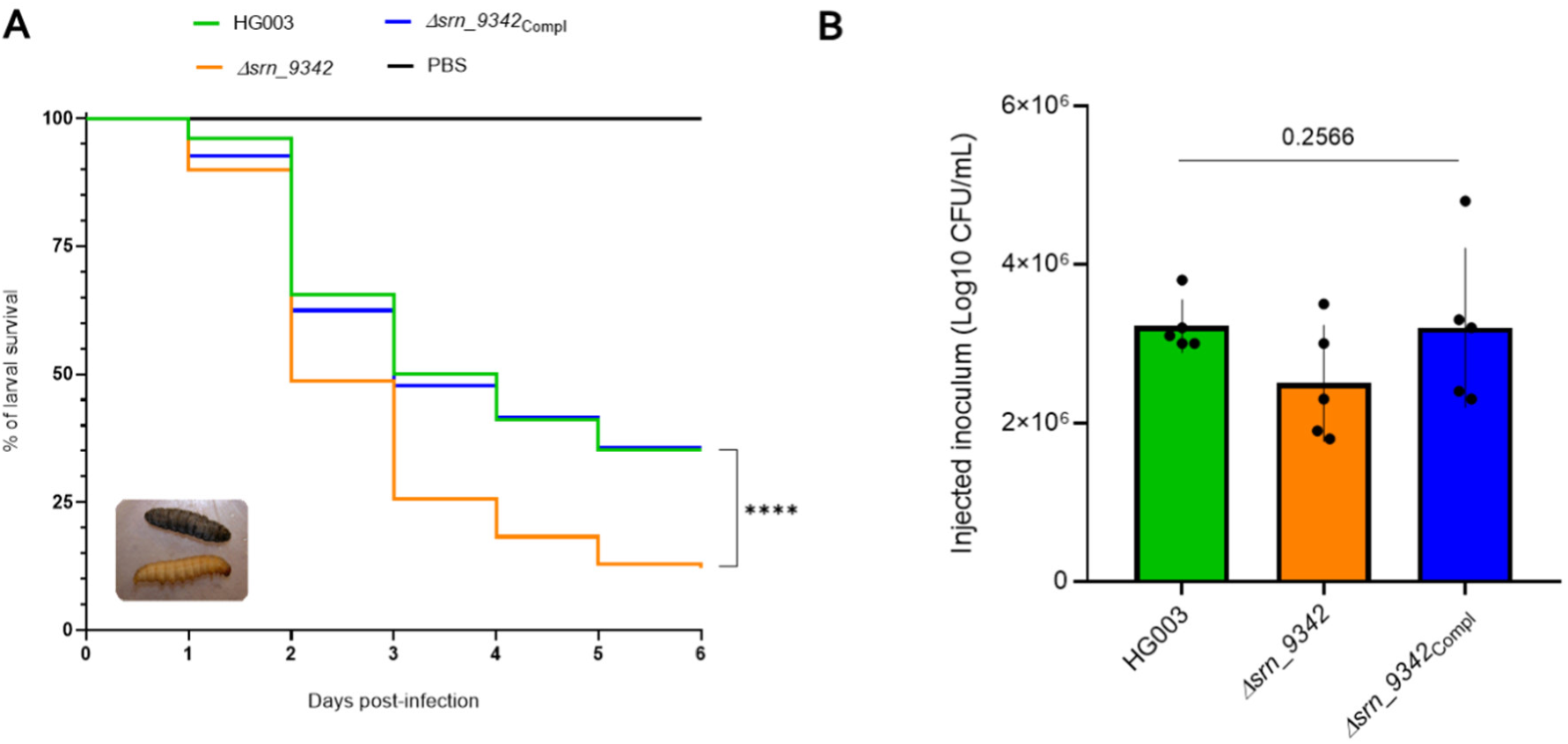
Srn_9342 is an anti-virulent factor in the *Galleria mellonella* model of infection. (A) Kaplan-Meier survival curves of larvae over six days post-infection with HG003, *Δsrn_9342* or *Δsrn_9342_Compl_* strains. PBS infected larvae was used as a non-infected viability control. Values are the mean of five biological replicates, with 148 larvae infected per strain in total. A Log-rank (Mantel-Cox) test was performed on the data (**** = p-value < 0.0001). (B) Verification of inoculum consistency across all replicates and strains. A one-way ANOVA test was performed on the data.

## DISCUSSION

sRNAs were initially defined in two categories based on their mode of biogenesis: *cis*-antisense encoded sRNAs or *trans*-encoded sRNAs. However, expansion of the knowledge on these regulators showed that it is way more complex with the discovery of sRNAs expressed from UTRs [38]. Here, we studied Srn_9342, an sRNA previously identified in *S. aureus* Newman strain and reported to be expressed under two forms (Srn_9342_S_ and Srn_9342_L_) of different length [20]. Recent study showed that the two forms are oppositely expressed as a function of growth phase and investigations on uncovering their targetome revealed that Srn_9342_S_ interacts with RNAIII, leading to an alteration of the expression of δ-hemolysin [21]. In the present study, we further characterized and investigated the role of Srn_9342 in the HG003 background, an NCTC_8325 derivative strain [23, 39]. HG003 appears as a reliable model to study virulence and stress responses as the functionality of *tcaR* (encoding a regulator of Protein A expression) and *rsbU* (SigB activator) genes were restored. In addition, the expression of RNAIII is more typical compared to Newman, in which it is overexpressed due to a mutation in the histidine kinase *saeS* gene [24].

First, we verified and confirmed that the expression profile of Srn_9342 in HG003 was similar than that reported in Newman (**Fig. 2**). Effectively, in this strain Srn_9342 is also expressed under two isoforms. Srn_9342_S_ and Srn_9342_L_ RNAs have distinct 3’ ends corresponding to putative intrinsic transcription terminators based on ARNold software predictions [20, 40]. This was corroborated by the analysis of the Srn_9342 expression profile in an HG001 *Δrho* mutant strain (**Fig. 3**) [31]. Similar to Newman, *srn_9342* gene overlaps with the 3’ end of an autolysin amidase domain gene (**Fig. 1**), an organization that is always retrieved except in the Mu50 strain where a second copy of the gene is present in an intergenic region [20]. This organization is typical of 3’ UTR-derived sRNAs [38] which are subdivided into two sub-categories based on the presence of an own promoter allowing transcription initiation (Type I), or whose expression results from the cleavage by a specific RNase of the mRNA expressed from the upstream gene (Type II) [16]. So far, type II 3’ UTR derived sRNAs are the most numerous, with for instance CpxQ sRNA produced from the RNase E cleavage of *cpxP* 3’ UTR mRNA in gram-negative bacteria [41], or RsaC released after RNase III cleavage of the 3’ UTR of *mntABC-rsaC* operon mRNA encoding a manganese ABC transporter in *S. aureus* [18]. To date, few type I sRNAs were described such as DapZ in *Salmonella typhimurium*, an sRNA that possesses its own promoter and shares the same Rho-independent transcription terminator of *dapB* mRNA [42].

To determine whether Srn_9342 is a type I or a type II 3’ UTR-derived sRNA, we studied its transcriptional activity using a fusion with *blaZ* encoding β-lactamase (**Fig. 3 and S2**). Cloning of the genic fragment located just upstream the coordinate corresponding to the 5’ end of Srn_9342 enabled reporter gene expression. Additionally, complementation of the *Δsrn_9342* mutant with its promoter at a neutral genomic locus (SAPhB1508) [34], fully restored the parental Srn_9342_S/L_ pattern of expression (**Fig. 5, S3, S4 and S5**). Together, these findings demonstrate that Srn_9342 is an independent transcriptional unit and therefore a type I 3’ UTR derived sRNA. To the best of our knowledge, this is the first RNA belonging to this class uncovered in *S. aureus*.

Study of reporter gene expression using β-lactamase and luciferase showed an increased level as a function of growth phase, suggesting an increased expression in a SigB-dependent manner (**Fig. 3 and 4**). In *S. aureus*, some sRNAs are reported to accumulate during post-exponential and stationary phase, and depend on the intervention of the SigB factor [43]. To determine whether this could be the case here, Srn_9342 isoforms expression was monitored by Northern blot in two SigB mutant, HG002 [23] and HG003 *ΔsigB* (this study, [33]). Whereas HG002 has a mutation in the SigB activator gene *rsbU*, HG003 *ΔsigB* is a knock out strain in which *sigB* is deleted. Srn_9342_L_ expression is drastically reduced in HG002 and null in HG003 *ΔsigB* whereas complementation with a plasmid expressing *sigB* restored the expression of Srn_9342_L_, confirming the crucial role of SigB for the biogenesis and/or stability of Srn_9342_L_, but not for Srn_9342_S_ (**Fig. 4**). Also, it indicated that the expression of the two isoforms may be independent although a complete understanding of the transition from the expression of Srn_9342_S_ to Srn_9342_L_ is not elucidated yet. Indeed, while reporter gene assays indicate an increase of gene expression, especially using firefly luciferase which should avoid reporter protein accumulation over time, quantification of the two isoforms by Northern blot indicated that the sum was somehow constant (**Fig. S1**). This suggests that the expression from a reporter system may not reflect the actual regulation of expression in the case of Srn_9342.

Even though the role of SigB is clear, Northern blot and transcriptional fusion do not demonstrate whether it binds the *srn_9342* promoter. Transcriptomic data collected by Mäder and colleagues combined with the search for a SigB consensus binding site suggested that Srn_9342 (named S810) is SigB-dependent [31]. The authors identified a putative SigB consensus binding site at nucleotides −52/-76 upstream the TSS of *srn_9342* identified by Bronsard and colleagues [20]. Although cases of consensus relatively distant from the TSS are described for other SigB targets in *S. aureus* [32], this predicted binding site is not the actual one as fusions made with a 50 nts promoter length is sufficient to allow reporter gene expression. Conversely, the manually-identified consensus described in Figure 4A appears as a better candidate, especially as it corresponds to typical −10 / −35 nts boxes while sharing most of the bases essential for SigB binding [32].

To qualify a sRNA as a regulator, it is mandatory to identify a target on which it has a positive or negative impact in terms of transcript or protein amount, or on which it acts as a sponge [44]. We previously identified several RNA targets of Srn_9342_S_ and Srn_9342_L_ by MAPS in Newman strain [21]. RNAs found enriched with one or the other of the baits were mostly not redundant, suggesting that they have independent biological role, in the same way as observed for their gene expression. Here, we focused on the transcript levels of various targets at three different cell density corresponding to various Srn_9342_S/L_ balances. *hemQ* mRNA appeared as an attractive target as its transcript level increased by five-fold in the *Δsrn_9342* mutant (**Fig. 6**). Using the gel retardation assay, it was confirmed that these two RNAs interact *in vitro* while the IntaRNA program predicted that the 5’ moiety of Srn_9342 (from nts 74 to 91) binds the mRNA within its coding sequence (from nts 383 to 402). This indicates that Srn_9342 do not follow a canonical pattern in term of mechanism of action. Indeed, sRNAs usually bind the RBS of target mRNAs to inhibit ribosome loading, or bind the 5’ UTR of the mRNA to promote structural changes and favour RBS accessibility [45]. Srn_9342 is not the unique case of a *S. aureus* sRNA binding a coding sequence. A recent study showed that SprC targets the coding region of the *czrB* mRNA, encoding a zinc efflux pump, thereby destabilizing it [46]. In the case of Srn_9342-*hemQ* complex formation, it remains to be determined whether it leads to the recruitment of a double-strand endonuclease such as RNase III or RNase Y [47], or promotes transcription attenuation of *hemQ* mRNA due to the formation of an upstream stem-loop structure leading to premature transcription termination. In *S. aureus*, this latter mechanism has only been documented for RNAI, which regulates the replication of plasmid pT181 [48]. However, a similar regulatory mechanism has been linked to the virulence-associated RNA RnaG which interacts and modulates the transcription of *icsA* mRNA in the pathogen *Shigella flexneri* [49].

The interaction between Srn_9342 and *hemQ* leads to a repression of HemQ production, as shown by Western blot (**Fig. 6**), consistent with transcripts level alteration in the *Δsrn_9342* mutant. HemQ is part of the heme biosynthesis pathway in *S. aureus* [36], and mutation of its encoding gene leads to the formation of the SCV phenotype [26], as well as for *hemA* and *hemB* genes [27, 50]. Based on our findings showing a repression of HemQ by Srn_9342, we investigated whether the overexpression of the sRNA could alter bacterial fitness. Indeed, strains overexpressing Srn_9342 exhibited an increased lag phase and produced tiny colonies on agar plates, similar to SCVs. Consistent with previous report showing that SCV phenotype could be compensated by supplementation with hemin [26], the addition of this porphyrin resulted in partial restoration of colony size (**Fig. 7**), confirming the link between Srn_9342 and HemQ and therefore its role in regulating the formation of SCVs. However, since complementation is only partial, Srn_9342 may regulate other targets implicated in SCV phenotype that remain to be discovered.

sRNAs span all the functions required for bacterial adaptation and optimal physiology. In *S. aureus*, some of them are defined as virulence regulators, such as the pathogenicity island regulatory sRNAs SprC (anti-virulent) and SprD (pro-virulent) [13, 14, 51]. In a recent study, Srn_9342_S_ was reported to hybridize specifically the 3’ end of RNAIII, modulating the amount of δ-hemolysin, similarly to SprY sRNA which was shown to indirectly regulate Rot and Ecb expression via its binding with RNAIII, leading to altered virulence [44]. Therefore, we used the *Galleria mellonella* invertebrate model of infection [37, 52, 53] to determine the role of Srn_9342 in virulence adjustment. Larval survival study clearly demonstrated a role of Srn_9342 in virulence, which is increased in the *Δsrn_9342* mutant, indicating that Srn_9342 in another example of anti-virulent sRNA (**Fig. 8**). Furthermore, the effect of Srn_9342 on the HemQ protein level which is known to be critical for optimal growth, energetic metabolism, survival and virulence in *S. aureus* may also participate to this phenomenon [50].

Altogether, this study led to the characterization of Srn_9342 in *S. aureus*, which exhibit some remarkable features. It is the first type I sRNA derived from a 3’ UTR in this species and should to be added to the list of SigB-dependent sRNAs. It exhibits an uncommon pattern of expression, with two independent isoforms differentially expressed as a function of growth. Finally, it is involved in virulence and SCV phenotype of this species, through direct regulation with RNAIII and *hemQ* mRNA.

## MATERIAL & METHODS

### Bacterial strains, growth conditions and cloning strategies

All strains used in this study are listed in Table S1. *Escherichia coli* strains, used as host strain for plasmid constructions, were grown in Luria-Bertani Broth (LB, Sigma) under agitation at 37 °C (except for mutant constructions with pIMAY plasmid) with 100 µg/mL of ampicillin. *S. aureus* was grown in Tryptone Soya Broth (TSB, Oxoid) under agitation (160 rpm) at 37 °C (except for mutant constructions with pIMAY plasmid) with chloramphenicol (10 µg/mL) or tetracycline (2 µg/mL) when necessary. 5 µM of Hemin (Sigma) was added to the medium to study SCV phenotype restoration. After an overnight preculture, *S. aureus* was diluted in fresh medium (+/- ATB) at an OD_600nm_ of 0.1 and bacterial growth was monitored by OD_600nm_ measurements at different time points. Plasmid generated were transformed by heat shock at 42°C into *E. coli* XL1-Blue and electroporated into *S. aureus* RN4220 then in *S. aureus* final strains; or into *E. coli* IM08B and electroporated directly in *S. aureus* final strains [54]. Cloning of pRIT-207nts plasmid was done with a Gibson Assembly Master Mix (New England Biolabs), which allows a one-step assembly of multiple PCR fragments into a vector. All others cloning were classically performed by vector and PCR products restriction for 2 hours at 37°C in Cutsmart Buffer (New England Biolabs) with appropriate restrictions enzymes (BamHI, EcoRI, KpnI, SacI, SmaI or SphI from New England Biolabs), and ligation done with T4 DNA ligase as recommended by the manufacturer (New England Biolabs). Table S2 lists plasmids used and constructed in this study. Table S3 lists the primers used in this study. Genetically modified strains HG003 *Δsrn_9342*, HG003 *Δsrn_9342*_Compl_ and HG003 *Δsrn_9342*_ComplShort_ were obtained by double homologous recombination using the temperature-sensitive vector pIMAY [55]. Complementation was performed in a neutral locus as described [34]. Inactivation of *sigB* gene in HG003 and HG003 *Δsrn_9342* was performed by phage α80 transduction [56] using a *S. aureus* WCUH29 *ΔsigB*::Tc mutant [33].

### Total RNA extraction

Bacterial pellets were obtained after centrifugation at 3500 rpm at 4°C for 10min. Cell pellets were resuspended in 500µL of lysis buffer (20 mM sodium acetate, 1 mM EDTA, 0.5% SDS, pH 5.5) and transferred into a FastPrep bead beater tube containing 250 µL of 0.1 mm glass beads (Sigma) and 500 µL of phenol (pH 4). Mechanical lysis was performed for 30 s at a power of 6.5 in a FastPrep-24 5G instrument (MP Biomedicals). Cellular debris were removed by centrifugation (10min, 4°C, max speed). Total RNA was isolated from the supernatant by phenol-chloroform extraction and ethanol precipitation as described [57]. RNA pellets were resuspended in an adequate volume of RNases free water (Eurobio) and quantified using a NanoDrop 2000 spectrophotometer (ThermoFisher Scientific).

### Northern blot analysis

Northern blots were performed using 10 µg of total RNA loaded and separated in 6% polyacrylamide / 8 M urea gels as previously described [58]. Hybond-N+ membranes (GE Healthcare) were hybridized with specific ^32^P-labeled probes (Table S3) in ExpressHyb solution (Clontech) for 2 hours at 37°C. The membrane was washed, exposed, and scanned with a Typhoon FLA 9500 phosphorimager (GE Healthcare). The image is acquired and processed using Image Quant TL software (GE Healthcare).

### RT-qPCR

For quantitative real-time PCR assay, 5 µg of total RNA were two-times treated with 5µL of Amplification Grade DNase I (Invitrogen, Carlsbad, USA) for 15 min at room temperature. A phenol-extraction was performed after each DNase treatments. cDNA synthesis was performed using a High-Capacity cDNA Archive Kit (Applied Biosystems) with random primers. Real-time PCR was done using a Power SYBR Green PCR Master mix (Applied Biosystems) using a StepOnePlus Real-Time PCR system (GE Healthcare). Primers used for specific qPCR are listed in Table S3. Data were analyzed using the comparative critical threshold (ΔΔCT) method as previously described: the target RNA amounts were compared with *gyrB* RNA, which served as internal control [14].

### Protein purification and Western blots

For total protein extracts, cell pellets were harvested form *S. aureus* TSB culture and resuspended into 100 µL of lysis buffer (10 mM Tris HCl pH 7.5, 1 mM EDTA, 0.1 mg/mL lysostaphin (Sigma)). Following incubation at 37°C for 15 min, 14µL of a 7X EDTA-free protease inhibitor cocktail (Roche) solution were added. A 5 sec sonication was performed on samples and protein quantity was measured using a Qubit and the Qubit protein assay kit (Invitrogen).

For Western blots, 5 µg of protein samples were diluted in Laemmli sample buffer (v1/1), denatured 5 min at 100°C, and separated on 16% Tricine-SDS-PAGE. In parallel, a second gel migration was similarly run but stained for 1 h with Ready Blue solution, then washed for 30 min with H_2_O to verify sample loading. Proteins were transferred onto Hybond-P PVDF membranes (Amersham). After overnight blocking at room temperature under agitation, membranes were incubated with 1:5000 anti-FLAG horseradish peroxidase-conjugated antibodies (HRP) (Sigma) for 1h. After several washes, membranes were revealed using an ECL Plus Western Blotting Detection Kit (Amersham), then scanned with an ImageQuant LAS 4000 imager (GE Healthcare).

### *In vitro* transcription and gel retardation assay

DNA template containing T7 promoter sequence upstream *hemQ* and *srn_9342* was generated by PCR using primers listed in Table S3. Purified PCR products were used as templates for *in vitro* transcription performed with the MEGAscript T7 kit (Ambion). RNAs were separated on a 6% polyacrylamide-7 M urea gel electrophoresis and eluted overnight in G50 elution buffer (20 mM Tris-HCl pH 7.5, NaCl 250mM, 1 mM EDTA, and 0.25% SDS). Then, RNAs were precipitated in cold ethanol 100% and 0.3 M of sodium acetate, and dephosphorylated using Quick CIP (New England Biolabs) according to the manufacturer’s protocol. RNA concentration was determined using Qubit and the Qubit High-sensitivity RNA assay kit (Invitrogen).

The 5′ end labelling of RNA was obtained with T4 polynucleotide kinase (New England Biolabs) and [γ^32^P] adenosine triphosphate as previously described [59]. Gel retardation assays were performed as previously described [60]. Briefly, electrophoretic mobility shift assays (EMSA) were performed with fragments of native RNAs (*hemQ*-496nts and Srn_9342-151nts) containing their predicted interaction domains. 3 fmoles of labelled *srn_9342-*151nts sRNA were incubated with increasing amount (from 162 to 6 000 fmoles) of unlabelled *hemQ* mRNAs, or with unlabelled 600 fmoles of Srn_9342-151nts and RsaD for specificity controls, then loaded on an 8% polyacrylamide gel under non-denaturing conditions. Gels were dried and visualized after 1 day of exposition using a Typhoon Phosphorimager (Molecular Dynamic). Data were quantified with ImageQuant software (GE Healthcare Life Science).

### β-lactamase and luciferase reporter assay

HG003 *Δsrn_9342* carrying pCN42-P*srn_9342*-*blaZ* fusions (different promoter lengths) were grown overnight under 160 rpm agitation at 37°C in TSB supplemented with 10 µg/mL chloramphenicol. Overnight cultures were diluted as described above and at time-points of interest, 2mL of culture were harvest and centrifuged 3 min at 13 000 rpm, 4°C. Pellets were resuspended in PBS 1X to obtain a cell density of 1 unit of OD_600nm_ per 100µL. Protein extracts were generated on 18µL of resuspended cells in which 2µL lysis buffer were added (Lysostaphin 1mg/mL, Benzonase (Milliport), MgCl_2_ 12.5 mM). Samples were incubated for 20 min at 37°C under agitation, then diluted at 1/2, 1/5 or 1/10 as a function of the signal intensity. Next, in three technical replicates per sample, 100µL of H_2_O, 2 µL of protein extract dilutions and 25µL of Nitrocefin (Sigma) 50 % (0.25 mg/mL) were placed in a 96-well microtiter plate and incubated in a Synergy 2 Multi-Mode Reader (BioTek) at 37°C under continuous shaking during 40 min. OD measurements at 492 nm were done every 10 min. After that, a Bradford protein assay was performed on protein extracts using BSA (100µg/mL) as a standard. In three technical replicates per sample, 100µL of H_2_O, 2.5 µL of 1/5 diluted protein extracts and 25µL of Bradford reactive (Sigma) were placed in a 96-well microtiter plate and incubated in a Synergy 2 Multi-Mode Reader (BioTek) at 37°C under continuous shaking. OD_595nm_ was measured after 30 sec. The average protein quantity standardized β-lactamase activity and standard deviation were calculated from three independent biological replicates. For luciferase reporter assay, strains caring pRIT-207nts plasmid were grown overnight under 160 rpm agitation at 37°C in TSB supplemented with 10 µg/mL chloramphenicol. To initiate kinetic, 60 µL of overnight cultures with addition of 3 µL of luciferin reagent were placed in 96-well black optically clear flat-bottom polystyrene plates (Revvity), with OD_600nm_ adjusted to 0.05, and incubated for 20 hours at 37°C under 160 rpm agitation in an Envision plate-reader (PerkinElmer). An automatic measurement of OD_600nm_ was performed every 10 minutes. RLU data was normalized based on OD_600nm_ at each time-point. Each independent experiment was performed with 4 or 5 technical replicates in the same plate.

### Study of S. aureus virulence in Galleria mellonella

*Galleria mellonella* larvae were raised in the laboratory under standardized conditions: under a 28°C incubation and fed with pollen balls and wax (La Ruche Roannaise, Besacier). Infections were performed on larvae weighing between 250 and 300 mg and not presenting phenotypic alterations (melanization, behavior) in order to standardize the trial. Selected larvae were incubated at room temperature for 1 night before injection, then disinfected externally with 70% ethanol. The infection was carried out using a KDS 100 automated syringe pump (KD Scientific) and a 1 mL syringe (Terumo) with a Microlance^TM^ 3 needle (BD). Haemocoel was injected with 10 μl of *S. aureus* strains at 10^6^ CFU/mL or 10µL of PBS for controls. The infected larvae were placed in square Petri dishes and incubated at 37°C. Mortality was monitored daily for 6 days, and larvae were considered dead when they were immobile, no longer responding to stimuli, and melanized. For each condition, five independent experiments were performed using 28 to 30 larvae for each infection, and 10 larvae for PBS control. Survival profiles were determined using the log-rank test and Kaplan-Meier survival plots.

