## Supplemental data for "A type I 3’ UTR-derived sRNA is involved in heme metabolism and virulence in *Staphylococcus aureus*"

**Table S1: Strains used and constructed for this study**

| Species | Strains | Characteristics | References |
|---|---|---|---|
| *Escherichia coli* | XL1-Blue | Derivative of *E. coli* K-12 strain : sgrR, guaC, crl+, lac−, supE, pgaA, narG, rssB, tonB, abgB, paaA, yncI, kdgR, rfbD, gatC+, gatB, gyrA96(nalR), luxS, recA1, rpoS, ygeY, endA1, ftsP, ttdB, glpR+, rph+, bglH, rbsR, thiE, cadB, cpdB, fimE, hsdR17 (rk−, mk+), λ− [F′::(yafJ-proBA-lacIqZΔM15-yaiL), Tn10(tetR)] | [61] |
| | IM08B | Derivative of DC10B strain with two sets of *S. aureus* CC8 *hsdMS* genes inserted at the chromosome (restriction modification system mimicking RN4220 *S. aureus* strain) | [54] |
| *Staphylococcus aureus* | RN4220 | NCTC8325 derivative used for transformations with plasmids constructed in *E. coli* XL1-Blue strain | [62] |
| | HG003 | NCTC8325 derivative strain restored for *rsbU* and *tcaR* | [23] |
| | HG001 | NCTC8325 derivative strain restored for *rsbU (tcaR -)* | [23] |
| | HG001 *Δrho* | HG001 deleted of *rho* gene | [31] |
| | HG002 | NCTC8325 derivative strain restored for *tcaR (rsbu -)* | [23] |
| | HG003 *ΔsigB* | HG003 deleted of *sigB* ($tet^R$) | This study |
| | HG003 *Δsrn_9342ΔsigB* | HG003 deleted of *srn_9342* and *sigB* ($tet^R$) | This study |
| | HG003 *Δsrn_9342* | HG003 deleted for 200nts promoter + *srn_9342* | This study |
| | HG003 $\Delta srn\_9342_{\mathit{Compl}\mathrm{Short}}$ | HG003 *Δsrn_9342* complemented in chromosomic SAPhB1502 neutral locus with 200 nts *srn_9342* promoter and *srn_9342* entire gene | This study |
| | HG003 $\Delta srn\_9342_{\mathrm{Compl}}$ | HG003 *Δsrn_9342* complemented in chromosomic SAPhB1502 neutral locus with 200 nts *srn_9342* promoter and *srn_9342* 160 first nts | This study |

### Table S2: Plasmids used and constructed for this study

| Plasmids | Characteristics | References |
|---|---|---|
| pCN41c | Plasmid containing *blaZ* reporter gene for transcriptionnal fusions derived from pCN41 : *ermC* (Erm$^R$) substituted by *cat194* (Chlo$^R$) in this study | [29, 30] |
| pCN41c-57nts | 50 nts upstream and 7 nts downstream *srn_9342* TSS in fusion with *blaZ* | This study |
| pCN41c-107nts | 100 nts upstream and 7 nts downstream *srn_9342* TSS in fusion with *blaZ* | This study |
| pCN41c-207nts | 200 nts upstream and 7 nts downstream *srn_9342* TSS in fusion with *blaZ* | This study |
| pCN35c | High-copy plasmid derived from pCN35 : *ermC* (Erm$^R$) substituted by *cat194* (Chlo$^R$) in this study | [29] |
| pCN35c P*rsbU-sigB* | pCN35c with *sigB* under *rsbU* promoter control | This study |
| pRIT | Plasmid containing *luc* reporter gene for traductionnal fusions (Chlo$^R$) | [63] |
| pRIT-207nts | pRIT with 200 nts upstream and 7 nts downstream *srn_9342* TSS in fusion with *sigA* RBS + *luc* | This study |
| pIMAY | Shuttle vector for chromosomal double-recombination event in *S. aureus* with *rep* thermosensitive | [55] |
| pIMAY-SAPhB1502 | pIMAY with homologies regions in SAPhB1502 and a EcoRI/EcoRV/SmaI MCS to insert sequences | This study |
| pIMAY-*srn_9342*$_S$ | pIMAY-SAPh1502 with 200nts *srn_9342* promoter + *srn_9342*160 first nts insert by EcoRI/SmaI plasmid restriction | This study |
| pIMAY-*srn_9342*$_L$ | pIMAY-SAPh1502 with 200nts promoter *srn_9342* + *srn_9342* insert by EcoRI/SmaI plasmid restriction | This study |
| pIMAY-*hemQ*-flag | pIMAY with homologies regions in *hemQ* and a 21aa flag sequence at *hemQ* 3' end for WB HemQ-flag protein detection | This study |
| pCN38 | Low-copy plasmid (Chlo$^R$) | [29] |
| pCN38 P*srn _9342*$_S$ | pCN38 with *srn_9342* 207nts promoter + *srn_9342*$_S$ | This study |
| pCN38 P*srn _9342*$_L$ | pCN38 with *srn_9342* 207nts promoter + *srn_9342*$_L$ | This study |
| pCN38 P*ami-srn_9342*$_L$ | pCN38 with *ami* promoter + *srn_9342*$_L$ | This study |

**Table S3: Primers and probes used for this study**

| Name | Characteristics | Utilization |
|---|---|---|
| NWMN_1141384 (Srn_9342) | GTAACGCACCTGCTTAAATAGACAT | Northern blots probes |
| tmRNA | ACACGCTTAATGAGCTCGGG | |
| HemQ F1 | GCCAGTGGTACCCGTTCGTCCTCTCCTTCAG | pIMAY for flag insertion at *hemQ* 3' end for Western Blot |
| HemQ R1 | TGTCATGATCTTTATAATCACCGTCATGGTCT TTGTAGTCAGAAATCGCAAAGAATTGATCG | |
| HemQ F2 | GGTGATTATAAAGATCATGACATCGACTACA AAGACGATGACAAGTAATACATTGGTACGTTT ATAAATTAATAAAAAAATTC | |
| HemQ R2 | GCCAGTGAGCTCATCGCAGTTAAAACATCAA TCAATC | |
| HemQ OUT FRW | CATAATCTGTTGCTTGTAATTGTG | |
| HemQ OUT REV | GTGTAACTTACGATCTAAAATCTCAG | |
| FRW in flag | CAAAGACCATGACGGTGATTATAAAG | |
| REV in flag | CATCGTCTTTGTAGTCGATGTC | |
| F HemQ Flag | GTATACGAAATGCGCTTTGATG | |
| R HemQ Flag | CTTATGTATACGGGAATATACTGAAG | |
| pIMAY_Forward | TACATGTCAAGAATAAACTGCCAAAGC | |
| pIMAY_Reverse | AATACCTGTGACGGAAGATCACTTCG | |
| qPCR_F_Srn9342 | AGGTATGTAACGCACCTGCT | RT-qPCR |
| qPCR_R_Srn9342 | TGGAATGGTTCTGCCCCACC | |
| HemQ qPCR FWD | CGCATTGCTGACTTCCTAATC | |
| HemQ qPCR REV | CTTGCTTTGATATGAGGGTTCTC | |
| Teg47 FWD | CGCAATCGCAATAACTGACAC | |
| Teg47 REV | TGTCCAACGCAATTAATATTGAGAC | |
| sprD FWD | TCAAGGAGCGCCTTTCATT | |
| sprD REV | TTGCGCTTTCCAAATCAATGT | |
| rsaC FWD | GCCATTCCCTACACACTCTTAT | |
| rsaC REV | CAGTTGTCCAATCACATAAGAACC | |
| SpxA FWD | GACTGAAGACGGTACTGATGAA | |
| SpxA REV | GACGACGTAATAAGCCAGGAT | |
| MvaS FWD | GGCGTCCAACTGGACATAAA | |
| MvaS REV | GGAAGCATAGAGATGCGAAGTC | |
| CidB FWD | ACTGGCGTCATGCTGAAT | |
| CidB REV | GATACCTACTGCGGCTGTTATAG | |
| HemQ qPCR FWD | CGCATTGCTGACTTCCTAATC | |
| HemQ qPCR REV | CTTGCTTTGATATGAGGGTTCTC | |
| SAOUHSC00699 FWD | CGCATTGCCGTTACTTTGATAC | |
| SAOUHSC00699 REV | CCTCAAGTAAGGTCGCCATTT | |
| RecA FWD | CGCCGAAGCATTTGTTAGAAG | |
| RecA REV | ACGTGAGTGTCTCCCATTTC | |
| FbaA FWD | GGTACTGTTGGTGGACAAGAAG | |

| | | |
|---|---|---|
| FbaA REV | ACCTAATGCTGGCGCTAATG | RT-qPCR |
| RNAIII FWD | AGATCACAGAGATGTGATGGAAA | |
| RNAIII REV | TCTTGTGCCATTGAAATCACTCC | |
| SirC FRW | GCACTAGGAATGAGTGGTTTAATG | |
| SirC REV | CACCACCTGTGATACCGATAAT | |
| vigR FRW | TTTCGAAATTCTCTGTGTTGGG | |
| vigR REV | GCTGGCCAATAGTTAGCTTTC | |
| SAOUHSC_2018_qPCR_F | TCTAGAACATCAAGTATTGGCAGT | |
| SAOUHSC_2018_qPCR_R | TCTTTGTTGTTTGGCTTTCGG | |
| GyrB qPCR FRW | GGTGGCGACTTTGATCTAGC | |
| GyrB qPCR REV | TAATATGCGCTCCATCCACA | |
| RACE_1141272_R1 | CTGATTAGGTGGGGCAGAACCATTCC | RACE mapping Srn_9342 |
| RACE_1141272_R2 | GAACCATTCCATGTTCTAATAGGCAAG | |
| RACE_1141272_F1 | CATAGTCATGTCTATTTAAGCAGGTGCG | |
| RACE_1141272_F2 (for Short) | GCAGGTGCGTTACATACCTGCTTTC | |
| RACE_1141471_F3 (for Long) | GATAACAGGCAGGTACTACGG | |
| M13 F | GTAAAACGACGGCCAGTG | |
| M13 R | CAGGAAACAGCTATGAC | |
| 5’ 9342 EMSA | GCCAGTTAATACGACTCACTATAGGGGGAGGGG CAACGTTATTA | *In vitro* Transcription (EMSA) |
| 3’ 9342S EMSA | GTAAATAGAAAGCAGGTATGTAAC | |
| 3’ 9342L EMSA | AAAAATAGGCAAGTACCGTAG | |
| 5’ HemQ496 EMSA | GCCAGTTAATACGACTCACTATAGG GATGAGTCAAGCAGCCGAAAC | |
| 3’ HemQ496 EMSA | GGCGTTCTTCCATAGTTAAC | |
| E_F2_9342_up | GGCTGGACTGACGGAATCG | Srn_9342 deletion in HG003 |
| F1_9342_up_KpnI | GGCTAGGTACCTTATGATTTCCCTATGTGG | |
| R4_delta_9342 | GGCAAGTACCGTTATGTATAATCACAATACCCAC CAGG | |
| F4_delta_9342 | GTATTGTGATTATACATAACGGTACTTG CCTATTTTTTTATG | |
| R1_9342_XmaI | TAGTGCCCGGGCCGCATCAGTTAAGATGC | |
| R3_delta_9342 | TTGTAAGTCATCAACTAACCTAC | |
| F_F3_9342_up | CTACATGGAAGAAAGTGCTAG | |
| R2_delta_9342 | CATTTGTAGTTATTTTAGCTTCAC | |
| Sequencing_9342delta | GTAACTCGCGGTCTATTGCC | |
| Locus3 P1 | GCCAGTGGTACCGGTCGAAAAAGCTAGAAAAGA AG | pIMAY for chromosomic complementation at locus SAPhB1502 with CDS EcoRI/EcoRV/SmaI |
| Locus3 P2 | GCCAGTCCCGGGGATATCGAATTCGGATAGAAA ACCAATCATCTTTATAGG | |
| Locus3 P3 | GCCAGTGAATTCGATATCCCCGGGAATAAAAAA GAAGAGAAGATGTAACACA | |
| Locus3 P4 | GCCAGTGAGCTCCTTCCTTGACTAATTGATCTGC C | |

| | | |
|---|---|---|
| Compl. Prom9342 FRW | GCCAGTGAATTCAAGTGCATGGAAAC GTAATAAATATG | pIMAY locus SAPhB1502 with *srn_9342*$_{S\ or\ L}$ and promoter *srn_9342* or *ami* |
| Compl.srn9342 S REV | GCCAGTCCCGGGTCTTTAAATGTAAATAGAAAGC AGGTA | |
| Compl.srn9342 L REV | GCCAGTCCCGGGATACTAAACAAATGTTTATTAG TAAAGTGT | |
| Compl.Pami FRW | GCCAGTGAATTCAAAATTTGTTTGATTTTTAATGG ATAATGTGATATAATGGGGGAGGGGCAACGTTAT TAC | |
| Locus3 OUT FRW | GCTAGAACAGCACTTGTTAACTTAG | |
| Locus3 OUT REV | ACAAATATAAACGGAGTTTGTTC | |
| pIMAY_Forward | TACATGTCAAGAATAAACTGCCAAAGC | |
| pIMAY_Reverse | AATACCTGTGACGGAAGATCACTTCG | |
| Srn_9342_F_SphI_HG003 | TCATATGCATGCAAGTGCATGGAAAC GTAATA | Overexpression of Srn_9342$_{S\ or\ L}$ under endogen promoter in pCN38/pCN35c |
| R9342S_BamHI_HG003 | GCCAGTGGATCCTCTTTAAATGTAAA TAGAAAGC | |
| R_9342L_BamHI_HG003 | GCCAGTGGATTCCAAATGTTTATTAG TAAAGTGT | |
| pCN35D | GTGCTGCAAGGCGATTAAGT | |
| pCN35G | GGCCTTTTGCTCACATGTTC | |
| Frw_prom_rsbU | CATATGCATGCACAAAATTGATAAGTGCAATTAA | pCN35c for SigB complementation (Prom*rsbU*-*srn_9342*) |
| Rev_prom_rsbU | ACTCTTTCGCCATCGATTAGTTGCCTCCTCACT | |
| Frw_sigB_rsbU | GGCAACTAATCGATGGCGAAAGAGTCGAAATC | |
| Rev_sigB | GCCAGTGGATCCACCCCTCGATTTCAATTTAA | |
| pCN35D | GTGCTGCAAGGCGATTAAGT | |
| pCN35G | GGCCTTTTGCTCACATGTTC | |
| Srn9342_F_pAMI_SpHI_ HG003 | TCATAGCATGCAAAATTTGTTTGATTTTTAAT GGATAATGTGATATAATGGGGGAGGGGCAA CGTTATTAC | Overexpression of Srn_9342$_{S\ or\ L}$ under constitutive *ami* promoter in pCN38/pCN35c |
| R9342S_BamHI_HG003 | GCCAGTGGATCCTCTTTAAATGTAAATAGAA AGC | |
| R_9342L_BamHI_HG003 | GCCAGTGGATTCCAAATGTTTATTAGTAAA GTGT | |
| pCN35D | GTGCTGCAAGGCGATTAAGT | |
| pCN35G | GGCCTTTTGCTCACATGTTC | |
| Prom50_9342_F | TCAGGGATCCTTATACAGAAGTGATGTTACAAGA TGGTCATGTTTGGGTAGC | Trancriptionnal fusions of Srn_9342 promoters in pCN41c (*blaZ*) |
| Prom100_9342_F | TCAGGGATCCTTCTTATCTTGTCCAGTGG GTTATC | |
| Prom150_9342_F | TCAGGGATCCCTAGATTCACAAACGGCAATC AACC | |
| Prom200_9342_HG003_F | TCAGGGATCCAAGTGCATGGAAACGTAATAAATA TGG | |
| Prom_9342_Rev _HG003 | TTCAGGAATTCCCCTCCCATGTATATCCTA | |
| BlaZ_Rev | TTGAAGCCTCAATAAGTGC | |
| pCN35G | GGCCTTTTGCTCACATGTTC | |

| | | |
|---|---|---|
| pRIT-P9342-F | CGGCATCAGAGCAGATTGTACTGAGAGAAGTGC ATGGAAACGTAATAAATATG | Trancriptionnal fusions of Srn_9342 207 nts promoter in pRIT (*luc*) |
| Luc-P9342+6 | GCCTTTCTTTATGTTTTTGGCGTCTTCCATGAAAC GGCCTCCCGACCCTCCCATGTATATCCTAC | |
| luc-F | ATGGAAGACGCCAAAAACATAAAGAAA | |
| pRIT-R | CTCTCAGTACAATCTGCTCTGATGCCG | |
| pRIT-F | CGGCATCAGAGCAGATTGTACTG | |
| seq from luc | AACGCGCCCAACACCGGCATAAAGAATTG | |

### SUPPLEMENTARY DATA

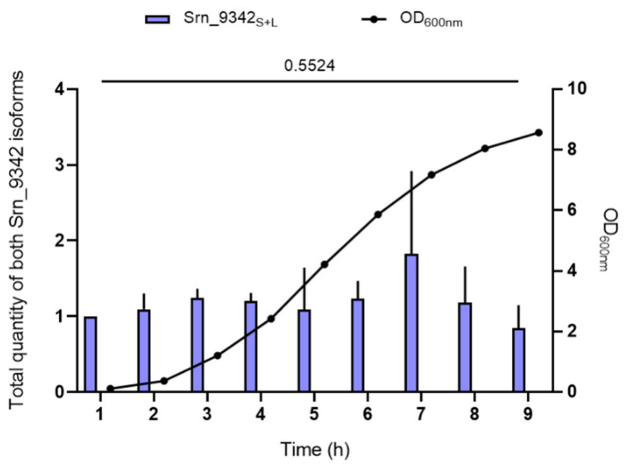

**Supplementary 1: The total level of both Srn_9342 isoforms is constant during growth.** Total quantity of both Srn_9342 isoforms (Srn_9342$_{S+L}$) at each hour of growth, standardized to the quantity at 1 hour. The Kruskal-Wallis non-parametric test was used for statistical analysis.

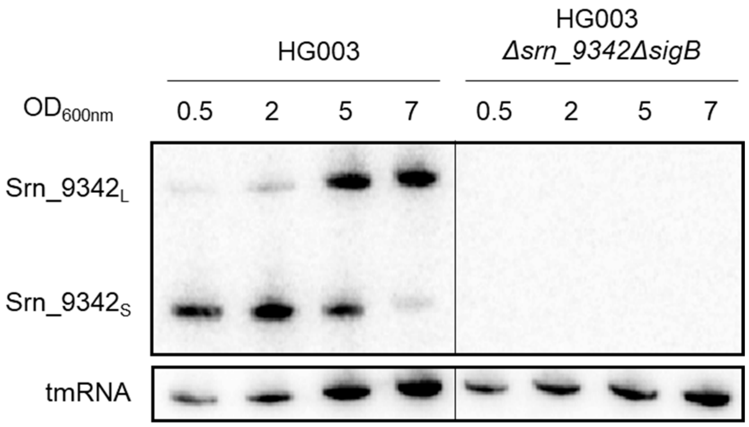

**Supplementary 2: Srn_9342$_S$ form detect in HG003 *ΔsigB* is not an artefact due to *ΔsigB* mutation phage transduction.** Northern blot of Srn_9342 isoforms in HG003 and HG003 *Δsrn_9342ΔsigB* during growth. tmRNA was used as an internal loading control. The Northern blot image is a representative experiment among three.

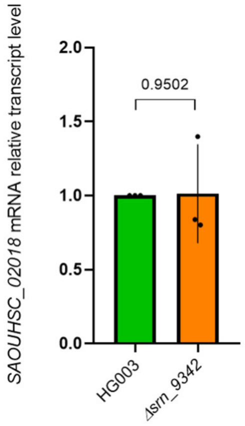

**Supplementary 3: Absence of polar effect of *srn_9342* deletion on the transcription level of the direct downstream gene *SAOUHSC_02018*.** Relative expression levels of *SAOUHSC_02018* mRNA determined by RT-qPCR on total RNA extracted from TSB cultures of HG003 and *Δsrn_9342* at $OD_{600nm}$ of 5. Values represent the mean of three biological replicates. Data were analysed using the ΔΔCT method, with *gyrB* as an internal control and the HG003 strain as the calibrator. An unpaired t-test was performed to determine significant differences among conditions.

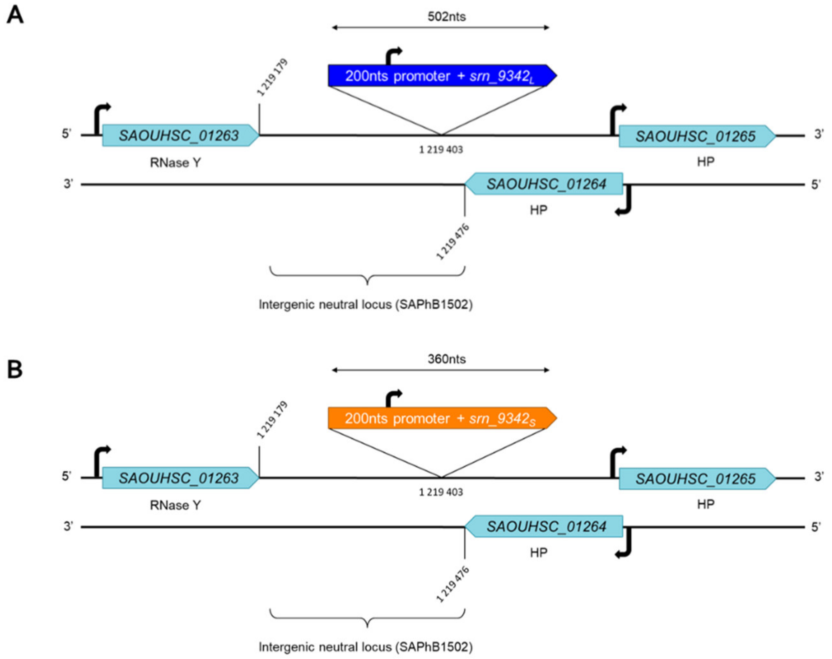

**Supplementary 4: Construction of Srn_9342 chromosomic complementation strains**. Insertion by double recombination of *srn_9342*$_L$ whole gene (A) or 160 nts *srn_9342*$_S$ gene (B) under control of 200nts *srn_9342* promoter in the SAPhB1502 neutral genomic locus of *Δsrn_9342* strain [34].

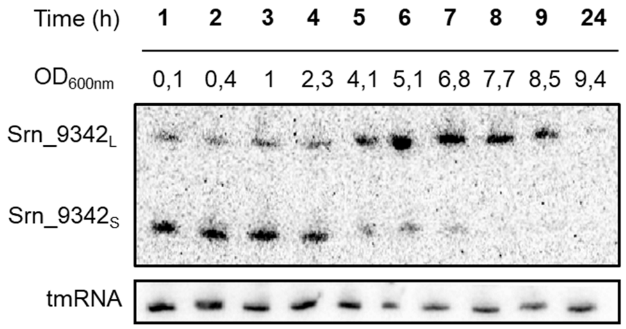

**Supplementary 5: Verification of Srn_9342 both isoforms expression in the strain HG003 *Δsrn_9342*$_{Compl}$.** Northern blot detection of Srn_9342$_S$ and Srn_9342$_L$ levels from total RNA extracted during TSB media growth of *Δsrn_9342*$_{Compl}$. tmRNA was used as an internal loading control. The Northern blot image is a representative experiment among three.

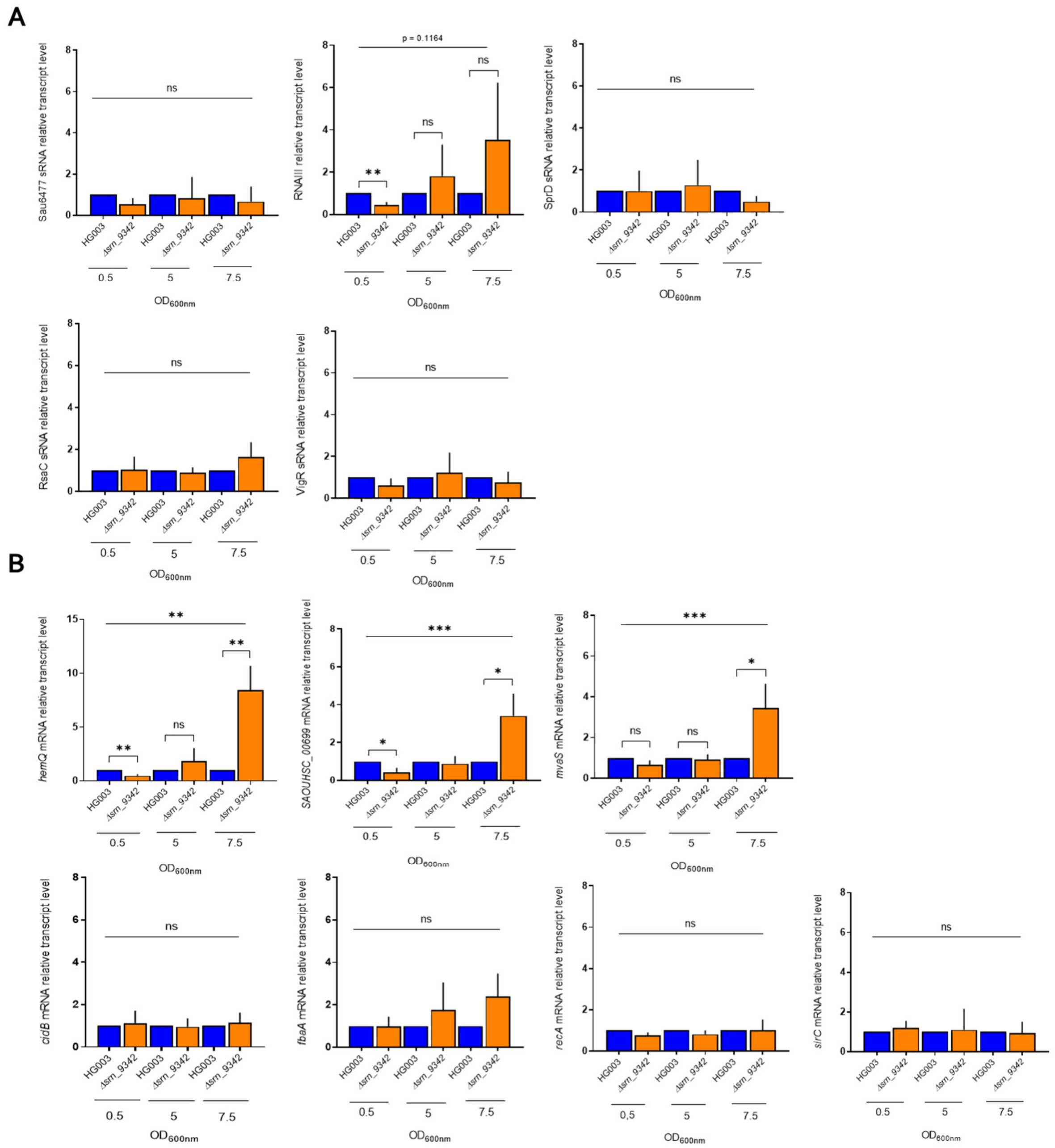

**Supplementary 6: Relative levels of putative RNA targets of Srn_9342$_S$ and Srn_9342$_L$ identified by MAPS during growth.** (A) Relative expression levels of 5 putative sRNA and (B) 7 putative mRNA targets (identified by MAPS) determined by RT-qPCR on total RNA extracted from TSB cultures of HG003 (blue) and *Δsrn_9342* (orange) at $OD_{600nm}$ of 0.5 or 5 or 7. Data were analysed using the ΔΔCT method, with *gyrB* as an internal control and HG003 strain as calibrator for each target. A One-way ANOVA followed by unpaired t-tests were performed to determine significant differences among conditions (ns = p > 0.05, * p < 0.05; ** p < 0.01; *** p < 0.001).

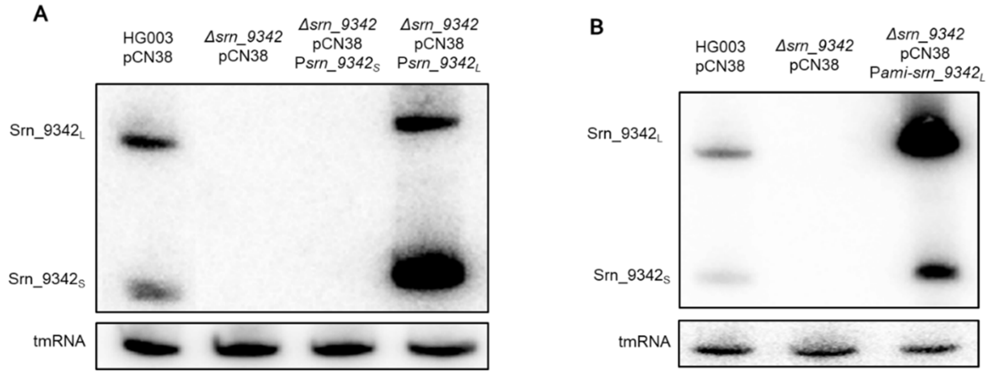

**Supplementary 7: Confirmation of Srn_9342$_{S+L}$ overexpression strains.** (A) Northern blot detection of Srn_9342$_S$ and Srn_9342$_L$ levels from total RNA extracted during TSB media growth of HG003 pCN38, *Δsrn_9342* pCN38, *Δsrn_9342* pCN38 P*srn_9342$_S$* (160 nts *srn_9342$_S$*) and *Δsrn_9342* pCN38 P*srn_9342$_L$* (full length *srn_9342*) at OD$_{600nm}$ of 5. tmRNA was used as an internal loading control. (B) Northern blot detection of Srn_9342$_S$ and Srn_9342$_L$ levels from total RNA extracted during TSB media growth of HG003 pCN38, *Δsrn_9342* pCN38 and *Δsrn_9342* pCN38 P*ami-srn_9342$_L$* (full length *srn_9342*) at OD$_{600nm}$ of 5. tmRNA was used as an internal loading control. *Δsrn_9342* pCN38 P*ami-srn_9342$_S$* (160 nts *srn_9342*) strain is not viable.
